# Adipose-driven complement-lipid reprogramming controls nociceptive vulnerability in obesity-associated osteoarthritis

**DOI:** 10.64898/2026.03.19.712962

**Authors:** Hope D. Welhaven, Joseph B. Lesnak, Kristin L. Lenz, Bethany Andoko, Arin K. Oesterich, Ashley N. Plumb, Katelyn E. Sadler, Luke E. Springer, Jacob G. Brockert, Darren Dumlao, Antonina M. Akk, Xiaobo Wu, Huimin Yan, Bo Zhang, Stephen P. Messier, Richard F. Loeser, Ronald K. June, John P. Atkinson, Christine T.N. Pham, Theodore J. Price, Farshid Guilak, Kelsey H. Collins

**Author notes:** Correspondence to: Kelsey H. Collins, Department of Orthopaedic Surgery, University of California San Francisco San Francisco, CA 94143.

## Abstract

Obesity amplifies osteoarthritis (OA) pain disproportionately to joint damage, creating a major unmet clinical need for non-opioid interventions that act beyond the joint. Using OA as a translational model, we integrated serum multi-omics in obese mice with surgically induced OA and genetic and adipose-reconstitution models of complement factor D (FD). In humans, we analyzed longitudinal metabolomics data from the IDEA weight-loss trial and conducted functional studies in dorsal root ganglion (DRG) neurons. Adipose-derived FD emerged as a regulator of systemic immunometabolic state: FD deficiency in obese mice worsened pain sensitivity whereas restoring circulating FD normalized pain and inflammatory markers without altering joint structure. Cross-species lipid profiling identified conserved shifts in linoleic acid versus arachidonic acid-derived lipids that were associated with pain phenotypes in mice and with pain improvement in humans. Defined lipid cocktails modulated excitability and TRPV1 sensitivity in human DRG neurons, and transcriptomics of knee-innervating DRGs revealed diet and FD-dependent activation of complement and neuronal excitability pathways. Together, these findings define an adipose-complement-lipid axis that regulates nociceptive vulnerability independent of joint damage and identify extra-articular targets for translational, non-opioid OA pain therapies.

**One Sentence Summary:** We identify an adipose-complement-lipid axis that systemically regulates sensory neuron sensitization, providing a mechanistic basis for pain-structure discordance in obesity-associated osteoarthritis.

## INTRODUCTION

Musculoskeletal diseases are leading contributors of chronic pain worldwide, and osteoarthritis (OA) is the most prevalent source of musculoskeletal pain. Clinically, pain – not radiographic joint damage – is the primary driver of disability, care-seeking behavior, and opioid use in OA (*1–3*). Yet pain severity in OA often diverges from the extent of structural joint damage, a discordance that is particularly pronounced in individuals with obesity (*4*). This clinical mismatch underscores a critical translational gap: the biological mechanisms by which obesity amplifies pain independent of joint damage remain poorly defined.

Obesity is one of the strongest risk factors for OA and powerfully amplifies the disconnect between joint pathology and pain. Obese individuals consistently report more severe pain for the same degree of structural damage across joints, including non-weight bearing joints such as the hand (*5*), compared to non-obese OA individuals, indicating that mechanical loading alone cannot explain obesity-associated OA pain. Even after weight loss, OA pain does not resolve (*6*), suggesting obesity induces persistent changes in pain-regulatory pathways and nociception. These clinical observations position OA in the context of obesity as a clinically relevant model to define how adipose-driven systemic signals modulate chronic pain independently of structural pathology, with direct implications for developing effective mechanism-based, non-opioid therapies.

Adipose tissue functions as an active endocrine and immunometabolic organ, secreting cytokines, complement factors, lipids, and metabolic mediators capable of reshaping inter-organ crosstalk and OA pathogenesis (*7, 8*). The complement system is a central innate immune pathway implicated in OA pathogenesis, yet prior work largely focused on excessive local complement activation within the joint as a pathological driver of tissue damage (*9, 10*). Far less is known about how dysregulated complement signaling – common in aging, obesity, and metabolic dysfunction – reprograms systemic mediators and engages extra-articular targets that influence nociception.

Complement factor D (FD), an adipose-derived serine protease (*11*) essential for alternative complement pathway (AP) activation, provides a direct mechanistic link between fat and OA pathology. Prior work demonstrated that adipose-derived FD regulates susceptibility to OA. In lipodystrophic (LD) mice lacking adipose tissue, protection from structural damage and pain is reversed by transplantation of healthy fat. Subsequent studies identified FD as a key adipose-derived mediators driving these effects, showing that loss of FD confers protection, whereas restoration of circulating FD derived from mouse embryonic fibroblasts (MEFs) reverses this phenotype (*11–13*). These findings position adipose-derived FD as a critical mediator of fat-joint crosstalk and an active driver of joint disease, rather than a passive bystander. Whether this axis operates similarly in obesity – the dominant clinical context in which OA develops – and whether it governs pain independently of joint damage remain unknown, with clear translational relevance for identifying new strategies to manage obesity-associated OA pain. While complement activity in OA is often viewed through the lens of local joint pathology, pain is the dominant clinical concern. The downstream targets of adipose-derived complement signals are undefined. It is unknown whether this axis operates primarily within the injured joint or reflects a systemic obesity-driven reprogramming that engages extra-articular targets, such as peripheral sensory neurons, to determine nociceptive vulnerability.

Here, we integrate serum multi-omics in obese mouse models of OA with longitudinal human metabolomics and functional assays in human dorsal root ganglion (DRG) neurons to define systemic mediators that couple adipose dysfunction to pain. We identify adipose-derived FD and alternative complement pathway signaling as obesity-modulated regulators of nociception that act largely independently of structural joint damage. Unexpectedly, FD deficiency reduced joint complement deposition yet exacerbated pain, revealing a dissociation between local complement activity and nociceptive outcomes and pointing to a dominant systemic regulatory role. Transcriptomic profiling of knee-innervating DRG neurons reveals HFD- and FD-dependent reprogramming of neuronal excitability pathways. Targeted lipidomics identified conserved shifts in circulating eicosanoid programs that tracked with pain phenotypes in mice and with longitudinal pain improvements in OA individuals. Guided by these signatures, we generated pain-promoting and pain-alleviating eicosanoid cocktails, which directly modulated nociceptor activity in human DRGs. Together, these findings reframe obesity-associated OA as a systemic immunometabolic disorder in which adipose-driven signals, rather than local joint pathology alone, determine nociceptive vulnerability. This framework opens new avenues for more effective therapeutic interventions to treat pain associated with obesity in OA.

## RESULTS

### HFD Exacerbates DMM-Induced OA and Shapes Systemic Complement-Lipid Signatures

A cohort of mice were fed either chow or HFD from weaning then challenged with destabilization of the medial meniscus (DMM) at 16 weeks of age and sacrificed at 12 weeks later to model obesity-associated OA. Both chow and HFD DMM mice exhibited greater joint damage, as shown by higher Modified Mankin and osteophyte scores compared to naïve controls; however, HFD further exacerbated structural damage (Fig. 1A-C), consistent with previous studies (*12, 14–16*). Synovitis increased in DMM limbs across both diets, but there were no significant differences between chow and HFD DMM animals (p=0.24) (Fig. 1D-E). HFD amplified pain-related measures, with pressure-pain thresholds demonstrating increased hyperalgesia in DMM limbs of HFD mice compared to chow-fed mice (Fig. 1F). These findings reinforce the role of obesity in worsening OA-related pain and structural damage, further shaping the joint microenvironment.

**Fig. 1.**
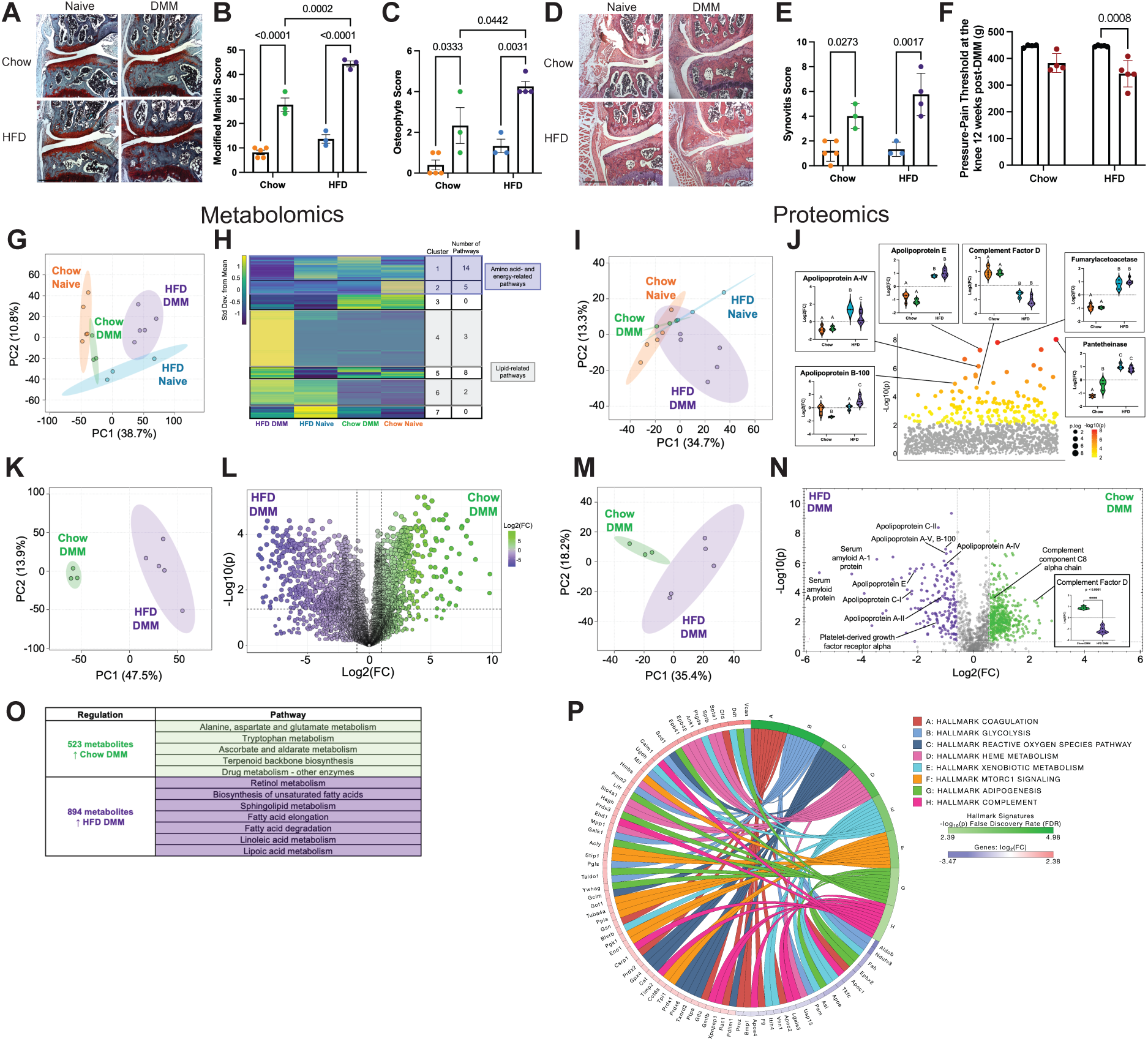
High-fat diet amplifies OA severity and reshapes systemic complement-lipid signatures. (A) Representative Safranin O/Fast Green-staining of the Medial Tibial plateaus from chow- and high fat diet (HFD)-fed mice naïve and DMM mice. Histological scoring including (B) Modified Mankin, (C) osteophyte, and (D) synovitis assessments with (E) representative H&E images where red indicates DMM animals and black indicates naïve animals. Scale bar, 400µm (F) Knee pressure-pain hyperalgesia measured by SMALGO in DMM (red) and contralateral limbs (black). (G-J) Serum multi-omic profiling across diet and injury groups. PCA analysis of (G) metabolites and (I) proteins. (H) Median metabolite intensity heatmap of ANOVA metabolites identifies clusters of metabolites for pathway analysis. (J) ANOVA identifies 185 differentially regulated proteins. To directly compare chow versus HFD DMM mice, metabolites were compared using (K) PCA and (L) volcano plot for (O) pathway analysis. Similarly, groups were compared at the protein level (M, N) revealing complement factor D is higher in chow DMM mice. (P) Pathway analysis using iPathway was performed, revealing complement and adipogenesis hallmark signatures. Orange – chow naïve, Green – chow DMM, Blue – HFD naïve, Purple – HFD DMM.

Previous work demonstrated that adipose-derived FD regulates OA susceptibility and pain (*11–13*), we next asked whether FD and related complement factors are reflected in systemic circulating profiles in the context of obesity and OA. To define these systemic changes, we performed unbiased proteomics and metabolomics on serum collected from DMM and naïve chow- and HFD-fed mice. A total of 8,532 metabolites and 1,243 proteins were detected across serum samples. Principal Component Analysis (PCA) showed clear separation between all four groups in both metabolomic (Fig. 1G) and proteomic analyses (Fig. 1I). Lipid-related pathways were significantly enriched in HFD DMM animals (Fig. 1H, Clusters 4 & 6), whereas amino acid and energy-related pathways were enriched in chow-fed groups (Fig. 1H, Clusters 1 & 2, Table S1). At the protein level, apolipoproteins such as A-IV and E were markedly higher in HFD mice, particularly in DMM animals (Fig. 1J). FD levels were lowest in HFD DMM mice and higher across chow-fed mice.

To further dissect systemic differences associated with diet in the context of OA, we performed PCA and volcano plot analysis comparing chow- and HFD-fed DMM mice (Fig. 1K,M). Volcano plot identified 894 metabolites enriched in HFD DMM mice compared to chow DMM controls, mapping to lipid pathways such as linoleic acid (LA), sphingolipid, and fatty acid metabolism (Fig. 1L,O). Conversely, chow DMM mice showed enrichment of amino acid pathways, ascorbate metabolism, and terpenoid biosynthesis. Proteomic analyses revealed significant upregulation of apolipoproteins (C-I, C-II, A-II, A-V, B-100, E) in HFD DMM mice, whereas FD and complement component C8 alpha chain were consistently highest in Chow DMM mice (Fig. 1N). Pathway analyses further highlighted enrichment of complement signaling and adipogenesis hallmark signatures (Fig. 1P, Table S2).

To delineate the contributions of obesity independent of injury-induced OA, we compared serum profiles of naïve chow- and HFD-fed mice. PCA analysis revealed clear separations between groups at both the metabolite and protein levels (Fig. S1 A,D), reflecting systemic reprogramming driven by diet diet-induced obesity. Volcano plot analysis identified 265 metabolites enriched in HFD naïve mice, mapping to lipid-related pathways including fatty acid and LA metabolism – patterns also observed in HFD DMM animals (Fig. S1B-C). In contrast, chow naïve mice exhibited enrichment of amino acid and central energy metabolic pathways. At the protein level, apolipoproteins (A-I, A-II, A-IV, E) were elevated in HFD naïve mice, whereas FD and complement component C8 (alpha, beta, and gamma) were highest in chow naïve mice (Fig. S1E). Pathway analyses confirmed consistent enrichment of complement signaling, regardless of injury status (Fig. S1F, Table S3). These findings indicate that obesity establishes a baseline systemic lipid and complement signature, upon which injury further reshapes the circulating environment to produce a distinct immunometabolic state in HFD DMM mice. Thus, circulating factors reflect both metabolic state and injury context, while further implicating obesity as a primary driver of the systemic changes associated with the OA phenotype.

### FD^−/−^ mice are not Protected from HFD-Induced OA Structure and Pain

To mechanistically define how adipose-derived complement signaling shapes obesity-associated OA, we next interrogated the role of FD, the obligate initiator and central gatekeeper of alternative complement pathway (*11, 13*). Although complement activation can be initiated through the classical, lectin, or alternative pathways, in mice the alternative complement pathway functions as a major amplification loop. Thus, genetic deletion of FD (FD^−/−^) selectively disrupts alternative pathway-mediated C3 and C5 production and substantially dampens downstream complement activation (*11, 17*). Therefore, we subjected wild-type (WT), FD^−/−^, and FD^−/−^ mice reconstituted with circulating FD via MEFs – a previously validated approach to restore systemic FD levels (*11–13*) – to HFD and DMM surgery (Fig. 2A). While previous studies have shown protection from DMM-induced OA on a chow diet (*13*), FD^−/−^ mice were not protected from OA structural damage or pain under HFD conditions. All three HFD-fed groups exhibited similar levels of joint damage, evidenced by Modified Mankin, osteophyte, and synovitis scores (Fig. 2B-F). MicroCT analysis of bone microstructure revealed no significant differences between DMM and contralateral limbs across groups (Fig. S2A-D). These findings suggest that FD deficiency does not mitigate structural OA outcomes in the context of obesity.

**Fig. 2.**
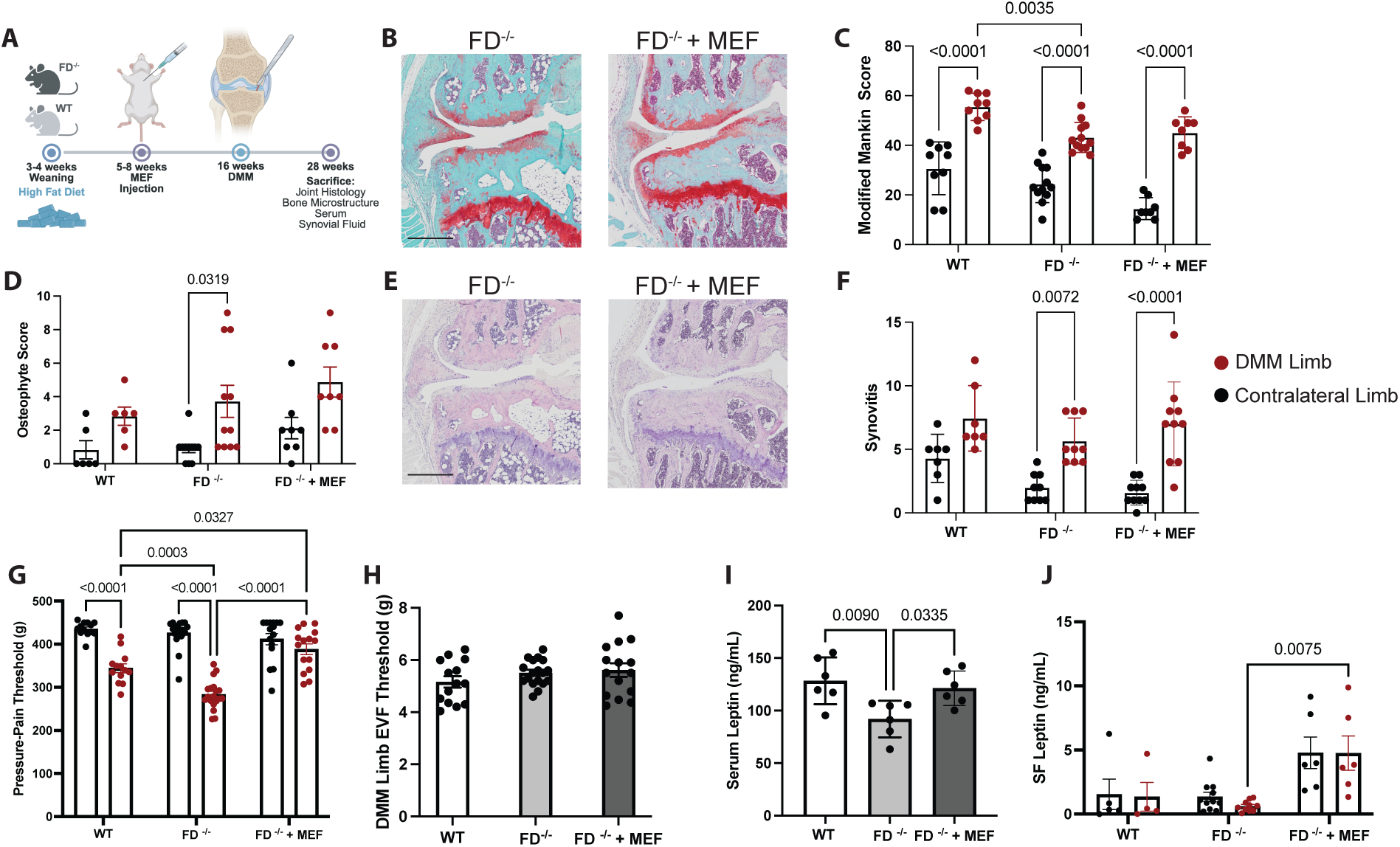
FD^−/−^ mice are susceptible to obesity-associated joint damage and pain. (A) Experimental timeline for HFD studies. (B) Representative Safranin-O/Fast Green and (E) H&E images with (C) Modified Mankin scores, (D) osteophyte score, and (F) synovitis scoring. Knee pain assessed by (G) SMALGO and (H) Electric Von Frey in DMM limbs. (I) Serum and (J) synovial fluid leptin. Data were analyzed using a 2-way repeated measures ANOVA with Tukey’s test for multiple comparisons (p<0.05). Black – contralateral limb, Red – DMM limb. Scale bar indicates 400 µm.

Pain phenotypes revealed distinct effects of FD. HFD FD^−/−^ mice demonstrated reduced pressure-pain hyperalgesia compared to WT, indicating increased hyperalgesia. In contrast, restoration of FD in FD^−/−^ + MEF mice increased pressure-pain thresholds relative to FD^−/−^ mice, consistent with reduced pain sensitivity (Fig. 2G). Notably, FD^−/−^ + MEF mice exhibited the highest pressure-pain thresholds across both limbs, indicating the lowest overall pain sensitivity among groups. Differences in allodynia between groups were not significant (Fig. 2H), collectively highlighting a disconnect between joint structural damage and pain sensitivity in FD^−/−^ mice under HFD. To further characterize adipose-derived systemic mediators associated with these FD-dependent differences in pain, we next examined circulating and synovial leptin levels. HFD FD^−/−^mice had reduced circulating leptin levels (Fig. 2I) compared to HFD WT and HFD FD^−/−^ + MEF. However, circulating leptin levels across all groups were still high (∼10x increase) compared to chow-fed groups previously observed (*13*). In synovial fluid, leptin levels were significantly higher in FD^−/−^ + MEF mice compared to FD^−/−^ mice (Fig. 2J).

Despite similar structural outcomes across groups, FD deficiency was associated with distinct pain phenotypes and marked alterations in systemic metabolic profiles. HFD FD^−/−^ mice gained weight similarly to HFD WT (Fig. S3A), but replenishing FD via MEFs mitigated weight gain and reduced total and abdominal fat (Fig. S3B-C), likely through repletion of C3 (*18*). Compared to chow-fed animals, HFD FD^−/−^ + MEF animals had increased MEF implant size, indicating the fat pad was functional and storing lipid (Fig. S3D). HFD FD^−/−^ mice demonstrated improved glucose tolerance compared to WT, but this protection was not reversed with FD reconstitution (Fig. S3E-F). Insulin resistance was heightened in HFD FD^−/−^ mice compared to WT and FD^−/−^ + MEF groups, which showed similar levels (Fig. S3G-H). FD deficiency was also associated with increased concentrations of circulating inflammatory mediators such as IL-1α, IL-6, MCP-1, and IL-17, which were normalized with FD reconstitution, whereas no differences were observed in circulating levels of IL-1ϕ3, IL-4, IL-10, or TNF-α (Fig. S4A-H). Apart from leptin, synovial fluid inflammatory profiles did not correlate with OA outcomes (Fig. S4I-L). Despite FD^−/−^not protecting the joints, across HFD-fed animals body fat correlated more tightly with abdominal fat (Fig. S4M) than overall body mass (Fig. S4N), reinforcing the systemic role of obesity in driving OA pathogenesis.

To explore if a complement bypass mechanism could explain the lack of structural protection in HFD FD^−/−^ mice, immunohistochemistry studies for C3 and C5-9 were conducted. Similar to chow-fed animals (*13*), HFD FD^−/−^ lacked detectable C3 (Fig. S5A-B) and C5-9 (Fig. S5E-F) in cartilage, suggesting that there is not a bypass mechanism conferring damage as well as heightened pain among FD^−/−^ mice. Even though FD repletion in HFD FD^−/−^ + MEF mice restored complement activation of C3 (Fig. S5C,D) and C5-9 (Fig. S5I,H) in cartilage, it did not mitigate joint damage as all HFD groups demonstrated similar levels of damage.

### DRG Transcriptomics Implicate Complement and Neuronal Pathways in Obesity-Associated OA Pain

To investigate the discordance between joint structure and pain in HFD FD^−/−^ mice and to explore the potential role of an adipose-nerve-pain signaling axis, we performed bulk transcriptomic analysis on L3-L5 DRGs – sensory neurons that innervate the knee joint and are responsible for transmitting pain signals to the brain. Comparative analyses between HFD WT and FD^−/−^ mice revealed distinct transcriptional profiles in DRGs associated with DMM-induced OA. Kyoto Encyclopedia of Genes and Genomes (KEGG) pathway analysis showed enrichment of complement and coagulation cascades in DRGs from HFD FD^−/−^ DMM limbs, while pathways such as IL-17 signaling, osteoclast differentiation, and histidine metabolism were enriched in HFD WT DMM limbs (Fig. 3A). Within HFD FD^−/−^ mice, transcriptional comparisons between DMM and contralateral limbs further highlighted pathways relevant to pain signaling. Glutamatergic synapse and calcium signaling pathways – both associated with neuronal excitability and pain sensitization – were enriched in DRGs from DMM limbs, alongside pathways such as long-term potentiation and amphetamine addiction (Fig. 3B). Contralateral limbs exhibited enrichment of amino acid metabolism pathways, including histidine, beta-alanine, arginine, and proline. These findings suggest that complement signaling, combined with neuronal excitability mechanisms, may drive the heightened pain sensitivity observed in HFD FD^−/−^ mice. Together, these data highlight complement and key neuronal pathways beyond the joint that contribute to pain in obesity-associated OA, reinforcing the systemic nature of disease pathogenesis and the role of complement as a mediator of adipose-nerve-pain signaling.

**Fig. 3.**
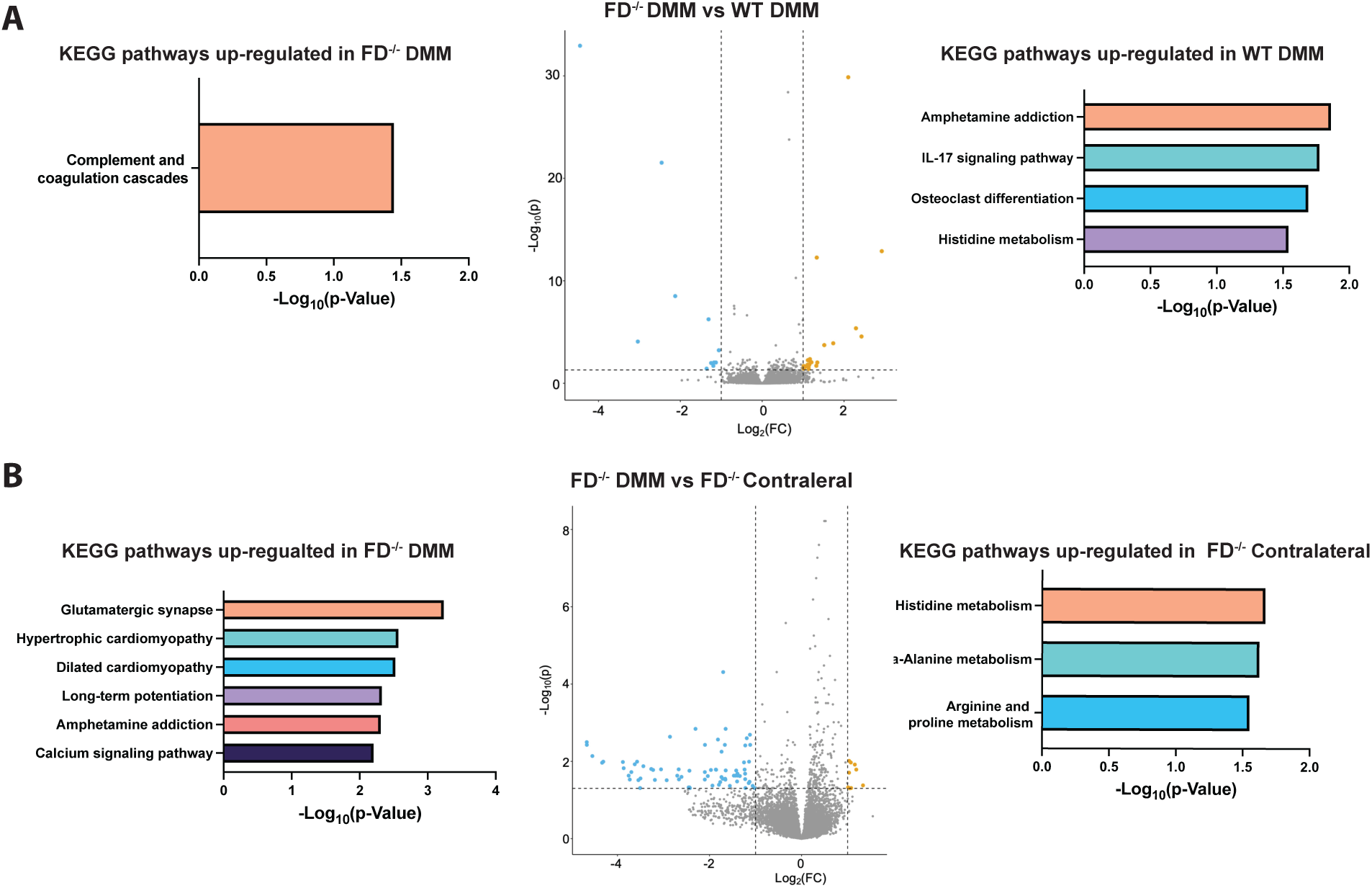
DRG neuron transcriptomic signatures reveal complement and neuronal pain pathway enrichment in obesity-associated OA. (A) Differentially enriched KEGG pathways in L3-L5 DRGs innervating DMM limbs of HFD FD^−/−^ (left) and HFD WT mice (right), with corresponding volcano plot (center). (B) KEGG pathway enrichment comparing DMM (left) and contralateral (left) DRGs in HFD FD^−/−^ mice, with corresponding volcano plot (center).

### Integrated Human and Mouse Analyses Identify Pain-Associated Eicosanoid Signatures

Distinct pain phenotypes observed in HFD-fed mice, alongside the detection of complement and lipid signatures as key metabolic nodes in WT naïve and DMM animals, prompted further investigation into systemic lipid and complement metabolism and their roles in pain modulation. Previously, our work revealed that loss of FD protects against cartilage damage following injury while driving acute pain and hyperalgesia, a process partly mediated by sex-specific alterations in eicosanoid profiles over time (*19*). Building on these findings of complement and eicosanoid dysregulation in OA pain, we sought to determine how FD deficiency and repletion in the context of HFD influence eicosanoid metabolism. PCA analysis showed partial separation among HFD-fed WT, FD^−/−^, and FD^−/−^+ MEF groups (Fig. 4A). ANOVA identified 22 significantly regulated eicosanoids – including eicosapentaenoic acid (EPA), 9-HODE, and docosahexaenoic acid (DHA) – that closely align with pain outcomes across groups (Fig. 4B). Targeted comparison of both groups of FD^−/−^ mice revealed clear group separation (Fig. 4C), highlighting distinct classes of systemic eicosanoids associated with pain outcomes. Eicosanoids elevated in FD^−/−^ mice were identified as candidate pain-promoting mediators (n=3) including 13, 14-dihydro-15-keto PGF2α; 5,6 DiHETE, and thromboxane B2 (Fig. 4D) while those enriched in FD^−/−^ + MEF mice were designated as candidate pain-alleviating mediators (n=10). This included 12,13 diHOME, LA, 12,13 EpOME, 9.10 EpOME, and others (Fig. 4E). These findings highlight FD repletion as a driver of systemic shifts favoring pain-alleviation, offering mechanistic insight into the partial protection from hyperalgesia observed in FD^−/−^ + MEF mice.

**Fig. 4.**
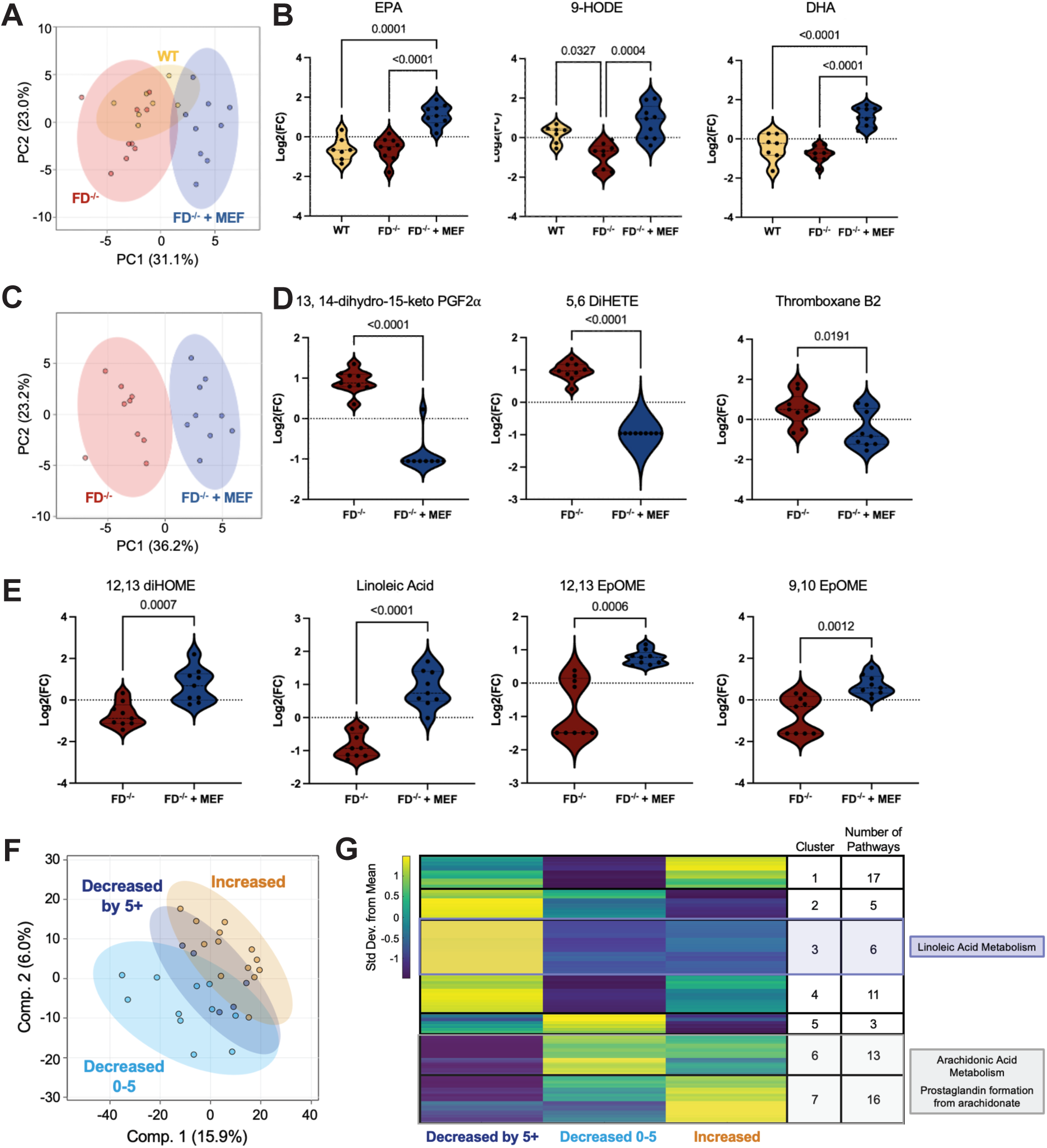
Conserved eicosanoid signatures link systemic metabolic state to OA pain across mice and humans. (A) Targeted serum eicosanoid profiling of HFD WT, FD^−/−^, and FD^−/−^ reveals clear separation of groups and (B) identified ANOVA metabolites that rend with pain phenotypes including EPA, 9-HODE, and DHA. (C) PCA comparing FD^−/−^ and FD^−/−^ + MEF. (D) Differentially regulated eicosanoids between groups where 13,14-dihydro-15-keto PGF2α, 5,6 DiHETE, and Thromboxane B2 were higher in FD^−/−^ mice. (E) Conversely, 12,13 DiHOME, linoleic acid, 12,13 EpOME, and 9.10 EpOME were higher in FD^−/−^ + MEF mice. (F) IDEA trial serum metabolomic profiles stratified by longitudinal change in WOMAC pain (increased, 0-5 decrease, >5 decrease). (G) Median metabolite intensity heatmap of metabolites identifies clusters of metabolites for pathway analysis highlighting eicosanoid pathways that are differentially regulated between groups, including enrichment of linoleic acid metabolism and arachidonic acid related pathways lowest in participants with the greatest pain reduction.

To explore whether these findings in mice parallel human OA pain outcomes, including in the context of obesity, we reanalyzed serum metabolomic data from a subset of Intensive Diet and Exercise for Arthritis (IDEA) trial participants, (previously published (*20*)). The IDEA trial evaluated the effects of 18 months of dietary-induced weight loss and exercise interventions on primary (knee joint compressive forces, IL-6 levels) and secondary outcomes (pain, function, mobility, quality of life) in obese individuals with knee OA (*21, 22*). Participants assigned to combined diet and exercise intervention exhibited the greatest weight loss, reductions in serum IL-6, and improvements in WOMAC pain and function scores (*21, 22*) as well as increased serum LA metabolism (*20*).

To model longitudinal dynamics, changes in metabolite intensity and WOMAC scores were calculated between baseline and 18 months. Participants were categorized into three pain trajectory groups: increased pain, decreased pain by 0-5 points, or decreased by > 5 points. Partial Least Squares-Discriminant Analysis (PLS-DA) was employed and revealed a partial separation of groups (Fig. 4F). Pathway analysis using a median metabolite intensity heatmap was employed to pinpoint clusters of similarly or differentially regulated metabolites on a global scale revealing conserved trends in eicosanoid metabolism (Fig. 4G, Table S4). Participants with the greatest pain reduction (> 5-point decrease in WOMAC pain scores) exhibited elevated LA metabolism, while arachidonic acid (AA) metabolism and prostaglandin formation were highest among those with minimal pain reduction (0–5 points) or worsening pain. This finding is consistent with the previously published data from the IDEA trial where LA metabolism was highest among diet and exercise individuals (*23*). It also aligns with findings from HFD-fed mice, where LA was associated with pain alleviation in FD^−/−^+ MEF mice.

LA is upstream of AA metabolism which triggers the generation and release of signaling molecules like prostaglandins. Both AA and LA belong to the family of bioactive eicosanoids and are lipid mediators that modulate inflammation and pain perception through cytokine signaling and COX2 activity (*24–26*). The conserved metabolic shifts observed in both mice and humans underscore the systemic role of eicosanoids in driving pain in obesity-associated OA. This convergence of species led us to test the functional roles of these eicosanoids in pain modulation.

### Functional Studies of DRG Neurons Demonstrate Pain-Promoting and Pain-Alleviating Functions of Eicosanoids Modulated by FD

Guided by these insights, composite eicosanoid “cocktails” enriched for pain-promoting or pain-alleviating mediators were generated based on mouse serum eicosanoid profiles (Table S5). Cultured human DRG neurons were treated with pain promoting or alleviating cocktails for 24 hours or 30 minutes and subsequently challenged with a low dose of capsaicin to test for TRPV1 sensitization (Table S6). Following 24-hour treatment, both the pain alleviation and pain promoting cocktails had a significantly lower proportion of capsaicin-responsive neurons compared to vehicle treated neurons (p.adj < 0.01), with no difference observed between the two cocktails (p.adj = 1.0) (Fig. 5A-C). No significant changes were found in the magnitude (F_2,264_ = 1.01, p=0.37), latency (F_2,264_ = 1.10, p=0.33), or AUC of the capsaicin response (F_2,264_ = 1.62, p=0.20) or the magnitude of the KCl response (F_2,283_ = 1.05, p=0.35) (Fig. 5D-G). However, there was a significant group effect for latency to KCl response (F_2,283_ = 12.19, p<0.01) as neurons treated with the pain promoting cocktail had a shorter latency to respond when compared to both vehicle (p<0.01) and alleviation cocktail (p<0.01) treated neurons (Fig. 5H). This data indicates that long-term treatment with both lipid cocktails decreases the total proportion of neurons responding to capsaicin, while the pain promoting cocktail alone reduces the latency of KCl evoked Ca^2+^ increases.

**Fig. 5.**
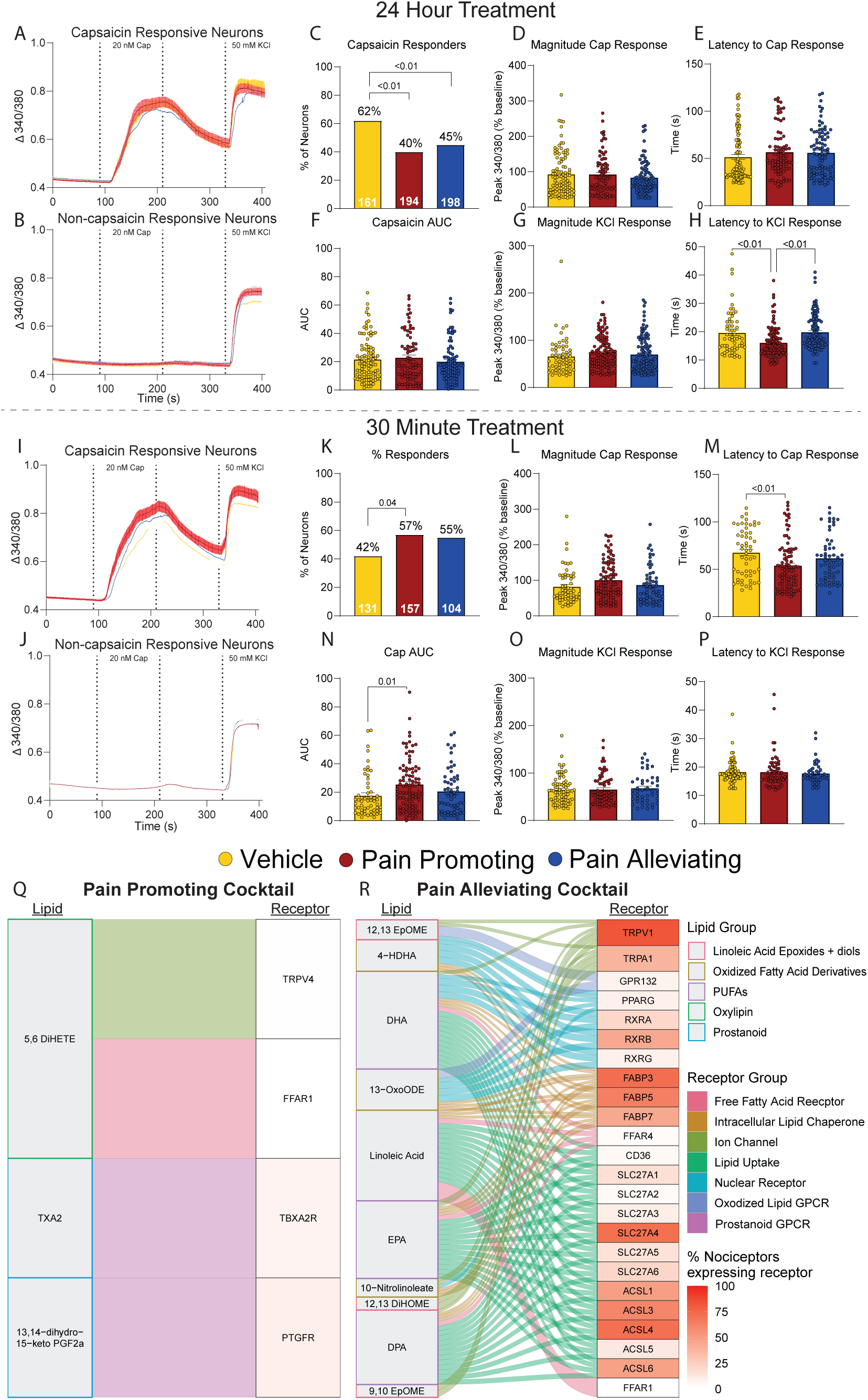
Pain promoting and alleviating lipid cocktails differentially modulate TRPV1 sensitivity in human DRG neurons. Cultured human DRG neurons were treated with a pain-promoting or alleviating lipid cocktail for 24 h or 30 minutes prior to challenge with low dose capsaicin during calcium imaging. (A-H) 24-hour pre-treatment. (A-C) Proportion of capsaicin-and non-capsaicin responsive neurons. Capsaicin response kinetics including (D) magnitude, (E) latency to respond and (F) AUC. KCl-evoked response including (G) magnitude and (H) latency to respond. (I-P) 30-minute pretreatment. (I-K) Proportion of capsaicin- and non-capsaicin responsive neurons with capsaicin response kinetics including (L) magnitude, (M) latency to respond, and (N) AUC. (O-P) No differences were observed in KCl-evoked responses. (Q-R) Lipid-receptor interactome illustrating possible interactions between cocktail components and receptors on human DRG neurons. Box color - lipid class; connection color - receptor type; receptor color - corresponds to percentage of human nociceptors previously identified to express this gene (*33*). Data is mean ± SEM.

In contrast, short-term (30 minutes) exposure to the pain promoting cocktail resulted in a significantly higher proportion of capsaicin-responsive neurons compared to the vehicle (p.adj = 0.04), while the alleviation cocktail had no significant effect (p.adj = 0.15) (Fig. 5I-K). There was also no significant difference in the proportion of capsaicin responsive neurons when comparing the two cocktails (p.adj = 1.0) (Fig. 5K). While there was no significant group effect for the magnitude of the capsaicin response (F_2,202_ = 2.48, p=0.09) (Fig. 5L), there was a significant effect for both the latency to respond (F_2,202_ = 5.36, p<0.01) (Fig. 5M) and AUC of the capsaicin response (F_2,202_ = 4.23, p=0.02) (Fig. 5N). Post-hoc analysis revealed that neurons from the pain promoting cocktail had a faster latency to respond (p<0.01) and higher AUC of the capsaicin response (p=0.01) when compared to vehicle treated neurons. There was no difference in the latency to respond (p=0.40) and AUC of the capsaicin response (p=0.60) between neurons treated with the pain alleviation cocktail and the vehicle. No group effects were observed for either magnitude (F_2,184_ = 0.18, p=0.83) or latency of KCl responses following short-term exposure (F_2,184_ = 0.24, p=0.79) (Fig. 5O, P). This data demonstrates that short-term exposure to the pain promoting cocktail sensitizes TRPV1 responses of human DRG neurons, evidenced by a higher proportion of capsaicin-responsive neurons, faster latency, and higher AUC of capsaicin-evoked responses. Finally, we explored if direct application of the lipid cocktail during calcium imaging resulted in acute increases in calcium in DRG neurons. Very little acute activation was demonstrated with either the pain promoting (5.2% of neurons) or alleviating cocktail (1.2% of neurons) which was not significantly different than vehicle treatment (1.4% of neurons) (Fig. S6). This absence of acute activation is likely due to the relatively low concentration of lipids in each cocktail.

To disentangle how each lipid ingredient could be altering human DRG neurons, we developed a lipid-to-human DRG neuron interactome. Components of the pain promoting cocktail were found to interact with potential receptors such as TRPV4, FFAR1, TBXA2R, and PTGFR (Fig. 5Q). The pain promoting cocktail resulted in TRPV1 sensitization following a short-term treatment but desensitization following long-term exposure. This could be occurring through activation of Thromboxane Prostanoid and Prostaglandin F receptors, which are G coupled GPCRs, which increase intracellular calcium and protein kinase C (PKC) (*27–30*) to sensitize TRPV1 channels in the short term (*31*) but lead to desensitization of the channel in the long term (*32*). There is likely little involvement of TRPV4 and FFAR1 due to their minimal expression on human DRG neurons (*33*).

In contrast, short-term treatment with the pain alleviation cocktail resulted in no effect on TRPV1 activity while long term treatment desensitized TRPV1 responses. The interactome revealed activation of several receptors located on human sensory neurons that could explain this effect including ion channels, cell membrane G-protein coupled receptors (GPCRs), and nuclear receptors (Fig. 5R). Several lipids in the pain alleviation cocktail – including 9,10 EpOME, 12, 13 EpOME, and 12,13 DiHOME – interact directly with the ion channels TRPV1 and TRPA1, which are known to be activated by lipids and could drive desensitization with prolonged activation (*34, 35*). Docosahexaenoic acid (DHA), eicosapentaenoic acid (EPA), docosapentaenoic acid (DPA) – all poly-unsaturated fatty acids (PUFAs) – interact with FFAR1/4, whose activity suppresses TRPV1 responses (*36–39*). Additionally, several lipids interact with nuclear receptors located in human DRG neurons such as CD36, SLC27/ACSL, and FABP3/5/7. 13-OxoODE and 10-Nitrolinoleate interact with the nuclear receptor peroxisome proliferator-activated receptors gamma (PPARγ) which forms a functional heterodimer with retinoid X receptors (RXR), including RXRA, RXRB, and RXRG (*40–42*). Thus, the pain alleviation cocktail could produce its TRPV1 desensitization effects through sustained activation of ion channels, FFARs, and nuclear receptors in human DRG neurons. Together, these findings demonstrate that eicosanoid signatures identified through cross-species lipid profiling are functionally active, with defined lipid combinations directly modulating human nociceptor excitability in a time-dependent manner.

## DISCUSSION

Obesity transforms OA from a localized joint disorder into a systemic immunometabolic condition in which pain severity is uncoupled from structural degeneration (*43*). Here, we identify adipose-derived complement factor D (FD) as a central regulator of this systemic state, linking alternative complement pathway balance to circulating lipid programs and nociceptive sensitization. Across mouse models and human cohorts, complement balance – rather than absolute complement activation – emerged as a key determinant of lipid class switching and pain outcomes independent of joint degeneration. Functionally, eicosanoid signatures derived from these systemic profiles directly modulated human nociceptor excitability, demonstrating that circulating lipid environments can act as pain-promoting or pain-alleviating signals. Together, these findings support a model in which adipose-driven complement-lipid signaling shapes nociceptive sensitization and underlies the discordance between pain and joint damage in obesity-associated OA.

Contrary to the canonical view of complement as a nociceptive driver, FD deficiency in HFD mice eliminated joint complement deposition yet exacerbated pain despite comparable structural damage. Restoration of circulating FD normalized pain, leptin, and inflammatory profiles despite reinstating joint complement detection. While prior work has largely focused on excess levels of complement factors (e.g., C3, C5) as mediators of nociception, our findings demonstrate that loss of complement activity can likewise promote pain. These findings indicate that in obesity-associated OA, FD does not primarily function as a local inflammatory amplifier but instead provides a systemic regulatory layer linking adipose biology to pain. This supports a model in which balanced alternative complement pathway signaling – not simply its magnitude – is required to maintain a pain-protective systemic state.

Because complement signaling is a pleiotropic pathway central to host defense and tissue homeostasis, making broad inhibition therapeutically challenging, we instead focused on downstream mediators through which adipose-derived signals engage nociceptive pathways. Integrated serum metabolomics across obese OA mice and human participants in the IDEA trial revealed conserved lipid signatures that tracked with pain trajectories rather than joint structure. FD deficiency, despite reduced joint complement deposition, favored pain-promoting eicosanoids such as 13, 14-dihydro-15-keto PGF2α, 5,6 DiHETE, and thromboxane B, whereas FD repletion restored a pain-alleviating landscape enriched in LA and cytochrome p450 LA-derived eicosanoids (DHA, EPA, DPA, 12,13 diHOME, 12,13 EpOME, 9,10 EpOME (*44*). These patterns were recapitulated in obese OA patients, where enriched LA metabolism associated with greater pain improvement, whereas AA metabolism and prostaglandin formation from arachidonate marked persistent or worsening pain. These concordant findings identify systemic lipid class switching as translationally relevant determinant of pain vulnerability in obesity-associated OA.

LA emerged as a central node whose downstream metabolites tracked with pain alleviation across species. Prior studies have linked LA to OA pathophysiology, including modulation of joint degeneration and synovitis (*16*), while AA, its downstream product, is proinflammatory (*45*), positively associated with obesity (*46*), detectable at higher levels acutely following knee injury in humans (*47*) and in early-stage OA synovial fluid (*48, 49*). Enrichment of omega-3 fatty acids (DHA, EPA) in FD-reconstituted mice with associated reduced pain is consistent with prior literature demonstrating that omega-3 fatty acids limit OA progression and promote tissue repair (*50, 51*), highlighting a conserved mechanism by which anti-inflammatory lipids constrain nociceptive sensitization. Our findings extend this framework by positioning lipid class switching as a systemic process that tracks with nociceptive outcomes across species.

Importantly, these lipid programs were not merely correlative. Pain-promoting and pain-alleviating cocktails derived directly from in vivo mouse and human signatures functionally modulated human DRG excitability. Short-term treatment with the pain-promoting cocktail sensitized DRG neurons to capsaicin, decreased latency to respond, and elevated AUC of the capsaicin-evoked signal. In contrast, long-term treatment with both cocktails reduced the total proportion of neurons responding to capsaicin, consistent with activity-dependent desensitization following sustained cocktail exposure, while the pain-promoting cocktail alone reduces the latency of KCl-evoked Ca^2+^ increases. These time-dependent effects demonstrate that circulating lipid environments may directly and dynamically modulate human nociceptor excitability rather than producing uniform pro- or anti-nociceptive effects.

To elucidate the mechanisms underlying these effects, we constructed a lipid-to-human DRG neuron interactome to identify receptor-ligand interactions that drive these responses. Pain promoting components were found to interact with TRPV4, FFAR1, TBXA2, and PTGFR, with the interactome analysis aligning with observed TRPV1 sensitization following short-term treatment and desensitization after long-term exposure. This is consistent with known mechanisms of Thromboxane, Prostanoid, and Prostaglandin F receptor activation, which increases intracellular PKC (*27–30*). PKC can sensitize TRPV1 channels in the short term (*31*), while prolonged activation may lead to desensitization (*32*). PKC activation may also explain the decreased latency of KCl-evoked Ca^2+^ increases observed with long-term treatment of the pain promoting cocktails, as PKC has been demonstrated to lead to increase neuronal excitability through modulation of Ca^2+^ channels (*52*). The eicosanoid 5,6-DiHETE interacts with TRPV4 but has been shown to be an antagonist of the TPRV4 channel (*53*), and TRPV4 shows minimal expression on human DRG neurons (*33*). Thus, this interaction likely does not contribute to the pain promoting cocktail TRPV1 effects. Finally, while FFAR1 activation suppresses TRPV1 activation in mouse DRG neurons(*39*), human sensory neurons express minimal FFAR1 (*33*).

The pain-alleviation cocktail exhibited time-dependent effects, with long-term treatment resulting in significant TRPV1 desensitization and short-term treatment showing no measurable impact on TRPV1 activity. These findings suggest that sustained exposure to the lipid components of the pain alleviation cocktail engages distinct receptor pathways to ultimately reduce TRPV1 activity. Insights from the interactome analysis identify several receptor families – including ion channels, GPCRs, and nuclear receptors – that may mediate these effects. For instance, persistent, low-level activation of TRPV1 by lipid components such as 9,10 EpOME, 12,13 EpOME, and 12,13 DiHOME can result in desensitization via dephosphorylation and internalization of the receptor (*54*). Additionally, PUFAs including DHA, EPA, and DPA interact with FFAR4, a receptor expressed on human sensory neurons that has been shown to suppress TRPV1 responses (*36–39*). The higher expression of FFAR4, relative to FFAR1 (*33*), further supports the role of receptor-mediated modulation in TRPV1 desensitization.

Nuclear receptor signaling may also contribute to the desensitization effects observed with prolonged exposure to the pain alleviation cocktail. Extracellular lipids can be taken up by cells through several mechanisms including passive diffusion (*55*). The cell surface fatty acid scavenger receptor CD36 (*56, 57*), and SLC27/ACSL uptake converts fatty acids into fatty acyl-CoAs to trap them intracellularly (*58, 59*). Once inside the cell, PUFAs and select fatty-acid derived lipids can be trafficked by cytosolic proteins, including FABP3/5/7, to the nucleus to interact with nuclear receptors (*60, 61*). Notably, 13-OxoODE and 10-Nitrolinileate interact with PPARγ receptors. While no direct *in vitro* rodent DRG work has been done on stimulation of PPARγ, activation of PPARγ has been shown to decrease neuropathic pain *in vivo* in pre-clinical animal models (*62, 63*). Thus, the pain alleviation cocktail could produce its TRPV1 desensitization effects through prolonged activation of ion channels, FFARs, and nuclear receptors in human DRG neurons. Collectively, these findings establish the DRG as a key extra-articular site at which systemic immunometabolic states are translated into nociceptive output. Taken together, in this framework sensory ganglia are not passive relays of joint-derived signals, but dynamic integration hubs that decode circulating immunometabolic cues.

Transcriptomic profiling of knee-innervating DRGs further revealed FD-dependent activation of complement-related and neuronal excitability pathways. While OA research has historically focused on joint tissues, emerging evidence implicates DRGs in sustaining chronic pain independent of structural damage. Prior studies demonstrate immune activation, transcriptional reprogramming, and macrophage infiltration with lumbar DRGs in experimental OA models (*64–66*). Our findings extend this paradigm by identifying complement- and pain-associated transcriptional signatures in knee-innervating DRGs and by demonstrating that circulating lipid programs directly modulate human sensory neuron excitability. Together, these data position periphery sensory ganglia as metabolic integration hubs in obesity-associated OA.

Viewed through a systems-level lens, these findings challenge the prevailing view that OA pain primarily arises from intra-articular pathology. Instead, we identify adipose-driven complement balance as a regulator of lipid class switching that directly tunes sensory neuron excitability, positioning the DRG as a critical extra-articular integration hub that translates systemic immunometabolic signals into nociceptive output. In this framework, the alternative complement pathway functions not simply as a local inflammatory amplifier within the joint, but as a systemic organizer of circulating lipid environments that shape pain vulnerability independent of structural degeneration. The dissociation between joint complement deposition and nociception underscores that metabolic imbalance can drive pain even when joint pathology is unchanged. By demonstrating that defined lipid profiles directly modulate human sensory neurons, we establish lipid class switching, and its downstream receptors, as a tractable therapeutic layer upstream of nociceptor sensitization.

Several limitations warrant consideration. Although we demonstrate functional effects of lipid profiles on human DRG excitability, the enzymatic intermediates linking complement balance to specific lipid pathways were not fully dissected. Additionally, experiments were performed in male mice. While our prior work demonstrated FD-dependent chondroprotection in both sexes and supports sexual dimorphism in FD in OA (*19*), future studies will define how sex-specific metabolic differences shape complement-eicosanoid-pain interactions within the context of obesity. Human metabolomic analyses were observational, limiting casual inference. However, concordant lipid signatures across mouse and human cohorts, together with functional validation in human neurons, support translational relevance.

Overall, our findings redefine obesity-associated OA pain as a disorder of systemic immunometabolic regulation rather than solely a direct readout of joint pathology. We demonstrate that adipose-driven complement balance orchestrates eicosanoid class switching that directly tunes nociceptor excitability, positioning the DRG as integration hubs that translate systemic state into pain vulnerability. In this framework, the alternative complement pathway functions not simply as an inflammatory amplifier within the joint, but as a systemic regulator of lipid class switching that constrains or promotes nociceptive sensitization independent of structural degeneration. These insights shift therapeutic strategy beyond cartilage preservation or intra-articular anti-inflammatory approaches. By identifying conserved circulating lipid signatures that stratify pain trajectories in humans and directly modulate human sensory neurons, our work establishes a mechanistic rationale for serum-based markers and extra-articular interventions targeting lipid pathways or their receptors. Rather than treating pain as a secondary consequence of joint damage, effective therapy in obesity-associated OA may require restoring immunometabolic balance. This systemic framework opens a tractable path toward effective strategies aimed at reducing nociceptive vulnerability at its metabolic source.

## MATERIALS AND METHODS

### Experimental Design

The objective of this study was to uncover mechanistic insights into the complicated role of alternative complement signaling in obesity-induced OA. In vivo experiments were conducted in adult male (16 to 28 weeks old) WT and FD^−/−^ mice to mirror a standard DMM trajectory model. Sample sizes were determined by power analyses (histology and pain testing: n=10/group, molecular assays: n=5/group, α=0.05). Surgeries were randomized and performed by one surgeon who was blinded to genotype. Investigators were blinded to diet, injury status, and genotype. No outliers were excluded. In vitro experiments used harvested DRGs from human donors (n=8). All procedures were approved by the relevant safety committees at University of California, San Francisco and Washington University.

### Animals and OA Induction by DMM

Male C57BL/6J mice were fed a HFD (60% kcal fat) or chow (10% kcal fat) from weaning. FD^−/−^ mice were bred as previously described (*11, 13, 17*). A subset of FD^−/−^ mice received a subcutaneous WT MEF implant to restore circulating FD (*13, 67, 68*). At 16 weeks, mice underwent unilateral DMM; contralateral limbs served as internal controls (*12, 13, 51, 69*). Mice were euthanized at 28 weeks. Body weight was monitored longitudinally; total fat mass was measured by DXA in FD cohorts.

### Joint Histology and Bone Microstructure

Knee joints were processed and scored using Modified Mankin, osteophyte, and synovitis scoring systems as previously described (*12, 13*). Safranin-O/Fast Green and H&E staining were performed using standard protocols. Subchondral and trabecular bone were analyzed by microcomputed tomography (Bruker SkyScan 1176) at an 18-um voxel resolution, consistent with previously reported methods (*12–15*). Main outcomes included bone mineral density (BMD) and bone volume fraction (BV/TV, %).

### Pain Assessments

Pain-related and behavior testing was performed one week prior to sacrifice (*12, 13*). Pressure-pain thresholds at the knee were measured using a Small Animal ALGOmeter (SMALGO) system (Bioseb) (*12, 15, 19*). Tactile allodynia was measured using Electronic Von Frey testing.

### Serum Multi-Omics

Serum from chow- and HFD-fed naïve and DMM mice was extracted and analyzed using a dual extraction protocol to simultaneously profiles proteins and metabolites (*13*). Proteomic analysis was performed by LC-MS/MS (timsTOF Pro2), and metabolomic analysis was performed using a SCIEX Exion 30 UPLC coupled to a SCIEX TripleTOF 6600+ at the UCSF Quantitative Metabolite Analysis Center. Data were processed using Spectronaut (proteomics) and MS Dial (metabolomic), followed by statistical and pathway analysis in MetaboAnalyst. Functional enrichment of differentially expressed proteins was performed using iPathways Guide.

### Targeted Eicosanoid Profiling

Serum from HFD WT, FD^−/−^, and FD^−/−^ + MEF mice was extracted to characterize systemic changes in eicosanoid profiles using our established targeted metabolomic panel of eicosanoids (*19*). Raw data was processed using a built-in SCIEX OS software package (version 2.1.6.59781) for peak picking, alignment, and quantitation. Eicosanoids detected were validated and confirmed with commercially bought standards.

### Serum and Synovial Fluid Profiling

A fasted blood sample was collected into BD microtainer SST serum separator tubes and allowed to clot at room temperature before serum isolation. SF was harvested using previously established methods (*12–15*). Both serum and SF were analyzed using Luminex multiplex 44-plex chemokine/cytokine assays (Eve Technologies) (*13*). Leptin levels in serum and SF were measured by ELISA (R&D Systems).

### Insulin and Glucose Tolerance Tests

Insulin and glucose tolerance tests were performed at 20 weeks following standard protocols (*12, 13*). Serial blood glucose levels were measured at 20-, 40-, 60-, and 120-minutes post-injection with a glucose meter (Contour; Bayer).

### Adipose and Dorsal Root Ganglion Bulk RNA-Seq

Bulk sequencing on L3-L5 DRGs was performed using our previously developed protocol (*13*). Libraries were prepared following ribosomal RNA depletion and sequenced on an Illumina NovaSeq platform. Reads were aligned to the Ensembl reference genome using STAR, and gene-level counts were generated with featureCounts. Differential expression and pathway analyses were performed using EnrichR.

### Intensive Diet and Exercise for Arthritis (IDEA) Trial Serum Metabolomic Profiling

Previously published serum metabolomics data from the IDEA trial (<u>NCT00381290</u>) were reanalyzed (*20–22*). Participants were stratified by longitudinal change in WOMAC pain scores over 18 months (increased, decreased 0-5, decreased >5 points). Metabolite changes were analyzed using MetaboAnalyst for multivariate and pathway analysis.

### Human DRG Recovery, Culture, and Calcium Imaging

Human DRGs were recovered from organ donors in Dallas TX through collaboration with the Southwest Transplant Alliance and under IRB-approved protocols. A total of 18 DRGs were used from 8 organ donors (Male n=5, Female=3, Table S7). As previously described (*70, 71*), human DRGs were surgically removed, dissociated, cultured, and maintained prior to treatment with pain-promoting or pain-alleviating lipid cocktails. For imaging, intracellular calcium responses were measured using Fura-2 (3ug/mL; Thermo Fisher Scientific, F1221). Neurons were stimulated with low-dose capsaicin followed by KCl to confirm viability. A ≥25% increase in the 340/380nm ratio was considered viable and included in the analysis. Detailed culture protocols, cocktail ingredient concentrations (Table S5), and calcium imaging parameters are detailed in the supplementary methods.

### Statistical Analysis

Inclusion criteria were established a priori based on successful completion of the experimental protocol, including survival through the study period and completion of surgical procedures and behavioral testing. No animals or data points were excluded from analysis. All animals meeting these criteria were included in downstream analyses. Data are presented as the mean +/− standard error. Statistical comparisons were performed using one- or two-way (group x limb, genotype x diet) ANOVA with Sidak or Tukey’s post hoc test. Assumptions of normality and equal variance were evaluated prior to parametric testing. For calcium imaging data, to compare proportion of capsaicin or lipid cocktail responsive neurons, a Fisher’s exact test was performed with a Bonferroni correction to adjust for multiple comparisons. For capsaicin and KCl kinetics analysis a one-way ANOVE with a Tukey’s post hoc test was used. Area under the curve (AUC) for the capsaicin evoked responses were calculated via the Simpson method. A priori α was defined as 0.05.

## List of Supplementary Materials

Materials and Methods

Fig. S1. Similar complement-lipid signatures are mirrored between naïve chow and high-fat diet.

Fig. S2. Bone microstructural assessments using microCT.

Fig. S3. Longitudinal analysis of body weight, fat, and glucose homeostasis.

Fig. S4. Metabolic and inflammatory dysregulation in HFD-fed mice.

Fig. S5. Assessment of complement activation in DMM joints using Immunohistochemistry (IHC).

Fig. S6. Cocktail application does not acutely activate DRG neurons.

Table S1. Differentially regulated pathways across chow- and HFD-fed DMM and naïve mice identified by median metabolite intensity heatmap analysis.

Table S2. Hallmark pathway enrichment analysis of differentially expressed serum proteins in HFD vs chow-fed DMM mice.

Table S3. Hallmark pathway enrichment analysis of differentially expressed serum proteins between HFD and chow-fed naïve mice.

Table S4. Differentially regulated pathways across IDEA participants by change in WOMAC pain scores over 18 months of weight loss intervention identified by median metabolite intensity heatmap analysis.

Table S5. Pain-promoting and pain-alleviating cocktails generated from HFD FD^−/−^ and MEF-treated HFD FD^−/−^ serum profiles and pressure-pain hyperalgesia outcomes.

Table S6. Statistical results for acute, short-, and long-term treatment of DRGs with pain-promoting and pain-alleviating cocktails.

Table S7. Organ donor demographic data.

## Acknowledgements

The original IDEA study was conducted under approval by the human subjects committee of Wake Forest Health Sciences (<u>NCT00381290</u>, registration data 10/2006). The authors thank the organ donors and their families for their gift and thank members of the Southwest Transplant Alliance for supporting the tissue recovery work.

## Author contributions

Conceptualization: HDW, JBL, KLL, AKO, CTNP, TJP, FG, KHC Methodology: HDW, JBL, ANP, AKO, XW, JPA, CTNP, TJP, FG, KHC

Investigation: HDW, JBL, KLL, BA, ANP, LES, JGB, DD, AMA, HY, BZ, XW, AHW, SPM, RFL, RKJ, JPA, AKO

Visualization: HDW, JBL, BA

Funding acquisition: JPA, CTNP, TJP, FG, KHC Project administration: HDW, JBL, TJP, FG, KHC Supervision: HDW, TJP, FG, KHC

Writing – original draft: HDW, JBL, BA, TJP, FG, KHC

Writing – review & editing: HDW, JBL, [add more once finished]

## Data and materials availability

All data supporting the findings of this study are included within the main text and/or the Supplemental Materials. Bulk RNA-seq data will be deposited in GEO. All other datasets will be deposited in Dryad and made publicly available upon publication. The FD^−/−^ mouse strain can be provided by JPA and XW pending review and completion of a material transfer agreement. Requests for such materials should be directed to JPA or XW.

