## Supplementary Materials for "Adipose-driven complement-lipid reprogramming controls nociceptive vulnerability in obesity-associated osteoarthritis"

### Materials and Methods

*Human DRG Recovery and Culturing*

Human DRGs were recovered from organ donors in Dallas TX through collaboration with the Southwest Transplant Alliance. All human tissue procedures are approved under protocol Legacy-MR-15-237 by the University of Texas at Dallas Institutional Review Board. Standard procedures and policies for donor screening and consent, and tissue recovery are handled by the Southwest Transplant Alliance and are approved by the United Network for Organ Sharing and the US Centers for Disease Control. Circulation of any donor medical information follows HIPAA regulation.

Human DRGs were surgically removed as previously described within 4 hours of cross-clamp(*69*). Upon recovery, DRGs were immediately placed in artificial cerebrospinal fluid (<1 hour) or Hibernate A (Fisher Scientific, NC0176976 ) supplemented with 1% N-2 (Thermo Scientific, 17502048), 2% NeuroCult SM1 (Stemcell technologies, 05711), 1% penicillin/streptomycin (Thermo Fisher Scientific, 15070063), 1% Glutamax (Thermo Scientific, 35050061), 2mM Sodium Pyruvate (Gibco, 11360-070), and 0.1% Bovine Serum Albumin (BSA; Biopharm, 71-040) (1-16hrs) at 4°C. Storage of full DRG bulbs in Hibernate A solution prior to dissociation results in DRG neuronal cultures with comparable electrophysiology and calcium imaging data as freshly dissociated neuronal cultures(*70*). A total of 18 DRGs were used from 8 organ donors (Male 5, Female 3). Demographic information is provided in Supplemental Table 7**.**

The bulb of thoracic and lumbar DRGs were exposed by trimming off excess connective tissue, fat, and nerve roots. The bulb of each DRG was diced into 3mm X 3mm sections and placed in 5mL of pre-warmed digestion enzyme consisting of 1 mg/mL of Stemxyme I (Worthington Biochemical, LS004106), 0.1 mg/mL of DNAse I (Worthington Biochemical, LS002139), and 10ng/mL of recombinant human β-NGF (R&D Systems, 256-GF) in HBSS without calcium and magnesium (Thermo Scientific, 14170-112). The tubes were placed in a shaking water bath at 37°C and the cell suspension was triturated every hour with fire polished

Pasteur pipettes until the DRGs were dissolved (4-5 hours). The dissociated tissue was filtered through a 100μM mesh strainer (Corning, 431752) and the resultant cell suspension was layered over 3mL of 10% BSA in HBSS in a 15mL tube. The tubes were centrifuged at 900g for 5 minutes at room temperature. The supernatant was aspirated and the pellet resuspended in prewarmed DRG media (BrainPhys® media (Stemcell technologies, 05790), containing 1% penicillin/streptomycin, 2% NeuroCult SM1, 1% Glutamax, 1% N-2, 10ng/mL β-NGF (R&D Systems, 256-GF-100), 2% HyClone™ Fetal Bovine Serum (Thermo Fisher Scientific, SH3008803IR), and 0.1% 5-Fluoro-2′-deoxyuridine (FRDU, Sigma-Aldrich, F0503). Cells were plated on 12mm coverslips pre-coated with 0.1mg/mL of poly-D-lysine (Sigma-Aldrich, P7405-5MG). The cultures were incubated at 37°C and 5% CO2 for 3 hours to allow for neuronal adherence. Following adherence, wells were flooded with prewarmed media, and half media changes were performed every other day.

*Human DRG Calcium Imaging*

Human DRG cultures were treated for 24 hours or 30 minutes with a pain promoting or alleviating lipid cocktail prior to calcium imaging (Table S5). For the 24-hour treatment, the lipid cocktails were prepared fresh the day of the treatment in serum starved media which consists of hDRG media lacking β-NGF, fetal bovine serum, and FRDU. The treatment was administered via a whole media change, and cultures were incubated at 37°C and 5% CO2 for 24 hours. For the 30-minute treatment, the lipid cocktails were prepared and delivered in the ENH calcium imaging bath solution (145mM NaCl, 3mM KCl, 2.5mM $CaCl^2$, 1.2mM $MgCl^2$, 10mM HEPES, 7mM Glucose, pH 7.4±0.1, osmolarity 320±3) prior to calcium imaging. In the acute treatment, the lipid cocktails were prepared and delivered in the ENH calcium imaging bath solution during data acquisition.

On the day of calcium imaging, coverslips were incubated with the calcium indicator Fura 2 (3ug/mL; Thermo Fisher Scientific, F1221) in ENH bath solution containing 2% BSA for 45 minutes at room temperature in the dark. Then to allow for desertification of the Fura dye the coverslip was incubated in ENH bath solution for 15 minutes at room temperature in the dark. For the 30-minute lipid cocktail treatment, the coverslips were exposed to the lipid cocktail during the desertification step which was extended to 30 minutes in length. Coverslips were mounted onto an inverted Nikon eclipse Ti2 microscope. Cells were perfused with ENH bath solution for 90 seconds, followed by stimulation with capsaicin (20nM, Sigma-Aldrich) for two minutes. Coverslips were perfused again with ENH bath solution for 2 minutes and then stimulated with 50mM KCl for 1 minute to confirm neuronal viability. For the acute treatment, the lipid cocktails or vehicle were applied for 5 minutes followed by a 2-minute ENH wash and then stimulation with 50mM KCl for 1 minute. During data acquisition images were captured every 500 ms at both 340 nm and 380 nm of excitation. For each neuron, the ratio of the 340/380nm fluorescent signal, with a background correction, was calculated using NIS Elements Software (Nikon, Version 5.42.03). A neuron was considered a responder to capsaicin or the lipid cocktail if it exhibited a ≥25% increase in the 340/380nm ratio when compared to the baseline recording. A neuron was considered viable and included in the analysis if it exhibited a ≥25% increase in 340/380nm ratio following KCl stimulation.

### Supplemental Figures & Tables

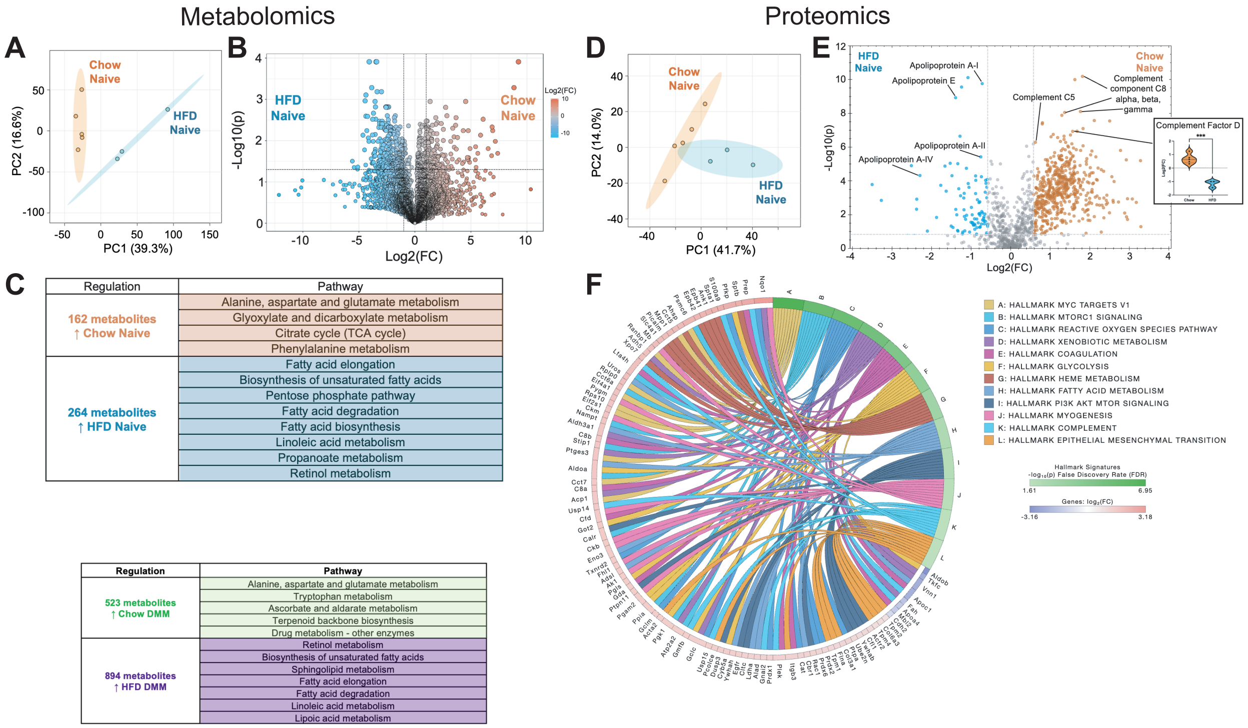


**Fig. S1. Similar complement-lipid signatures are mirrored between naïve chow and high-fat diet.** At the metabolite level, naïve animals – chow vs HFD – were compared using (A) PCA and (B) volcano plot for (C) pathway analysis. Similarly, groups were compared at the protein level (D, E) revealing complement factor D is higher in chow DMM mice. (F) Pathway analysis using iPathway was performed, revealing a complement signature.

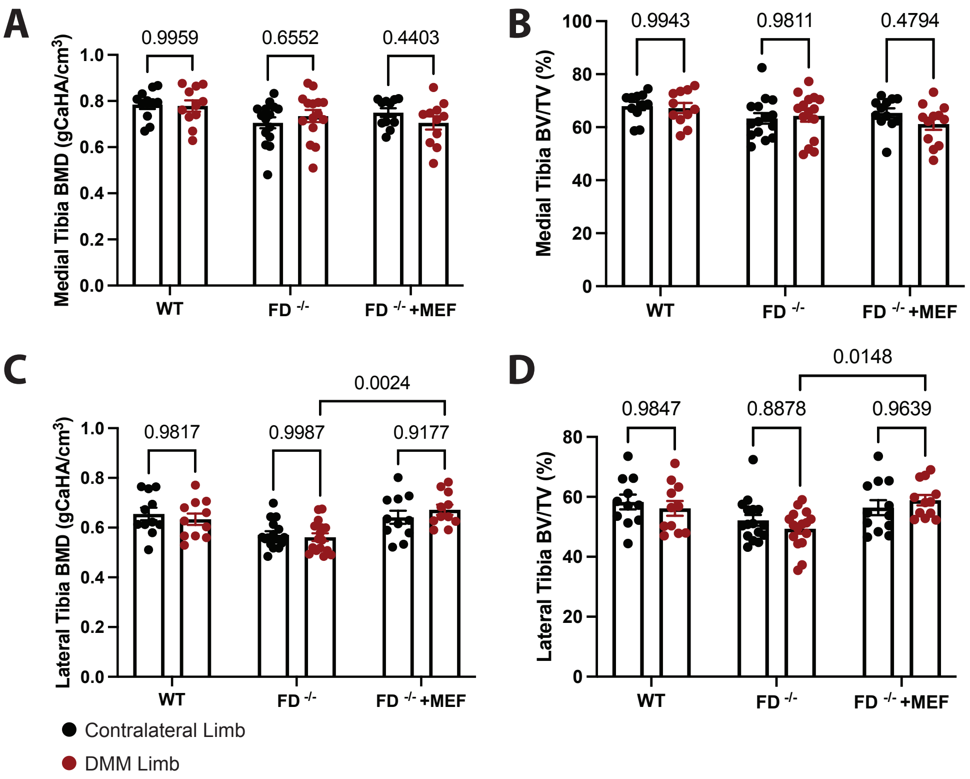


**Fig. S2. Bone microstructural assessments using microCT.** (A-B) Medial and (C-D) lateral tibia bone mineral density (BMD) and bone volume/total volume (BT/TV). Black – contralateral limb, Red – DMM limb.

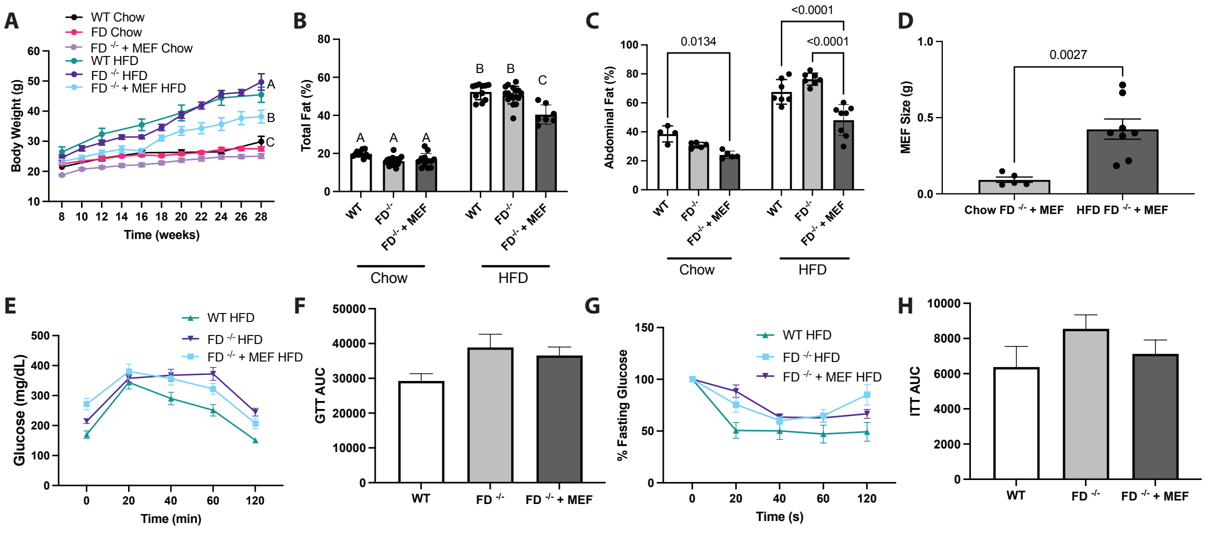


**Fig. S3. Longitudinal analysis of body weight, fat, and glucose homeostasis.** (A) Body weight over time, (B) total fat percentage, (C) abdominal fat percentage, (D) MEF implant size, (E) glucose tolerance test and (F) AUC, and (G) fasting glucose from insulin tolerance and (H) AUC.

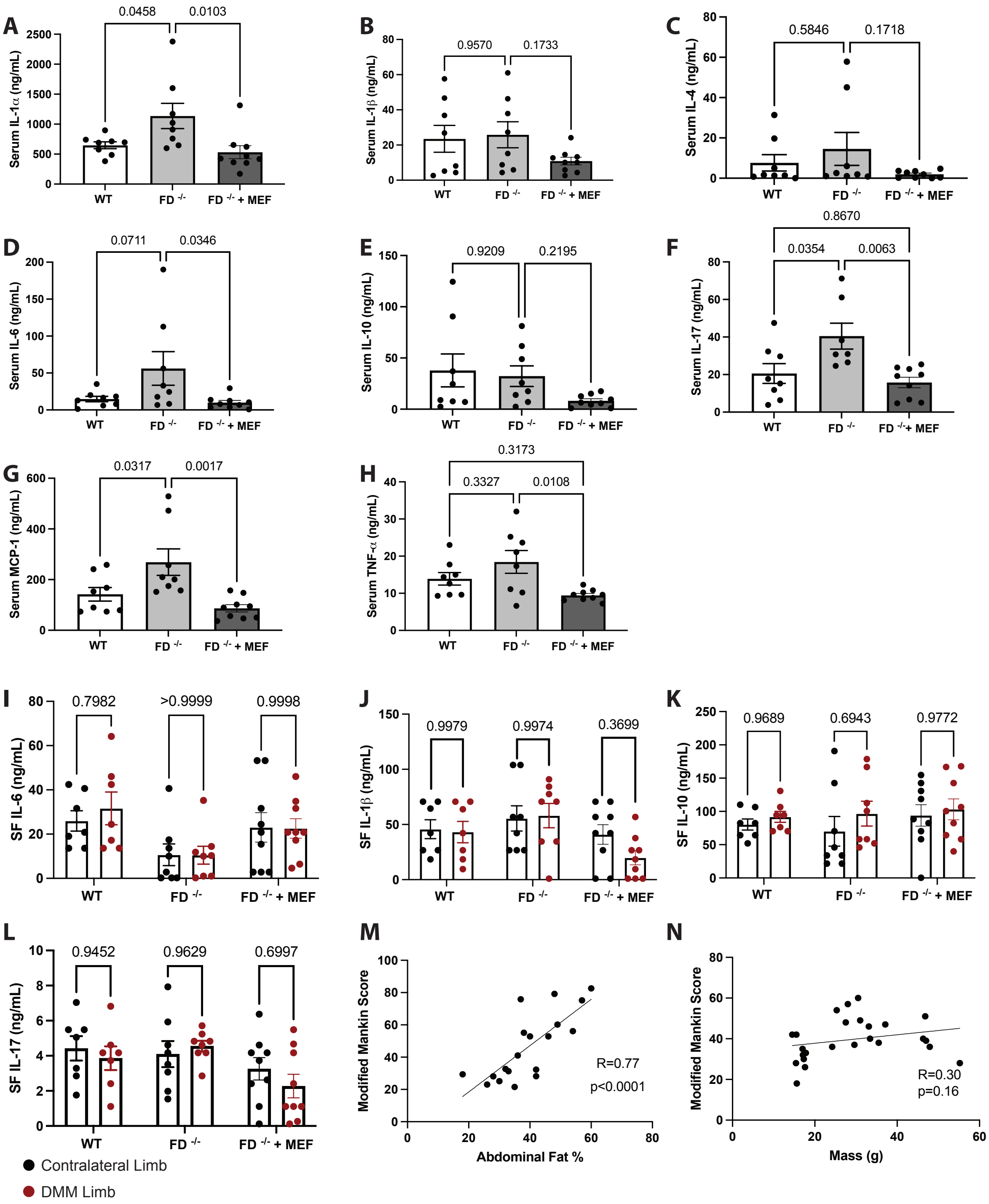


**Fig. S4. Metabolic and inflammatory dysregulation in HFD-fed mice.** (A) Circulating IL-1α, (B) IL-1β, (C) IL-4, (D) IL-6, (E) IL-10, (F) IL-17, (G) MCP-1, (H) and TNF-α. (I) Synovial fluid IL-

6, (J) IL-1β, (K) IL-10, (L) and IL-17. Spearman correlation between (M) abdominal fat and (N) body mass with Modified Mankin Score. Raw data are shown, n=7-14/group, one-way or two-way ANOVA with Tukey's or Sidak's post-hoc test, $p<0.05$.

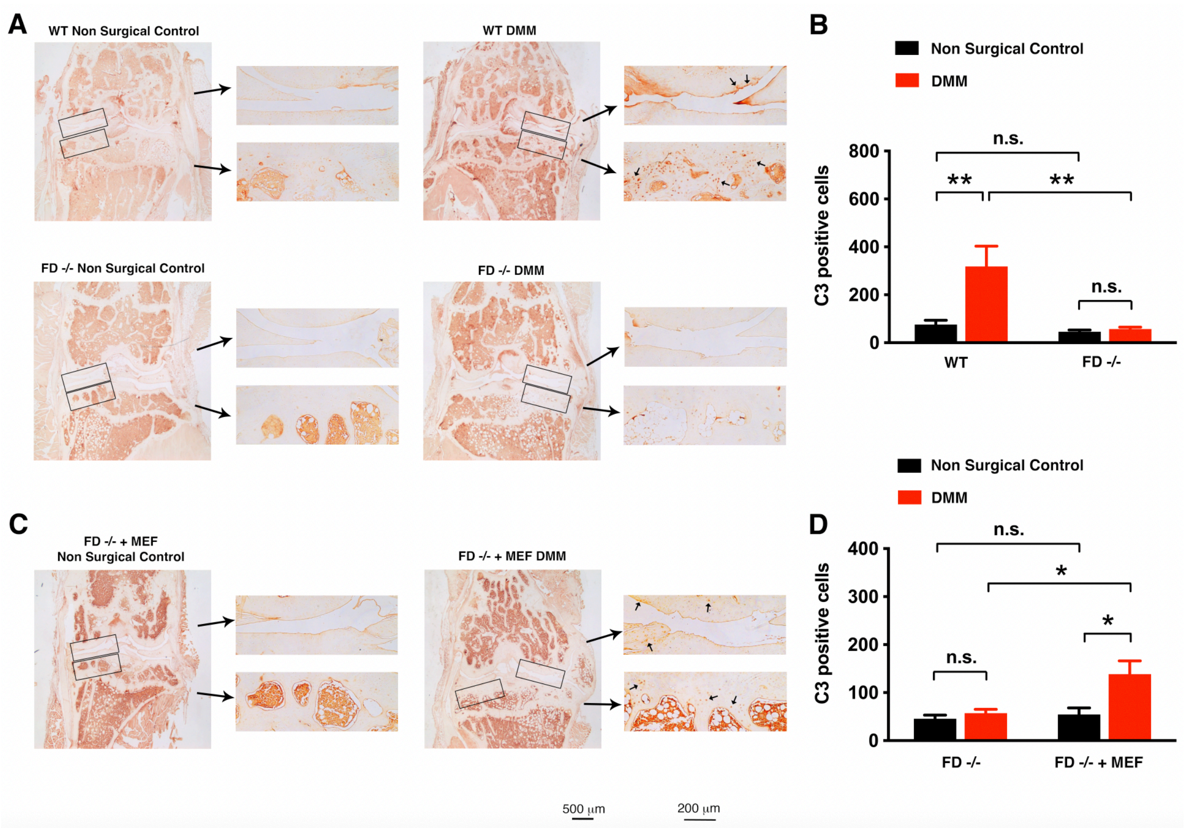

A
WT Non Surgical Control
WT DMM
FD -/- Non Surgical Control
FD -/- DMM
B
Non Surgical Control
DMM
C3 positive cells
800
600
400
200
0
n.s.
**
**
n.s.
WT
FD -/-
C
FD -/- + MEF
Non Surgical Control
FD -/- + MEF DMM
D
Non Surgical Control
DMM
C3 positive cells
400
300
200
100
0
n.s.
*
*
n.s.
FD -/-
FD -/- + MEF
500 μm
200 μm


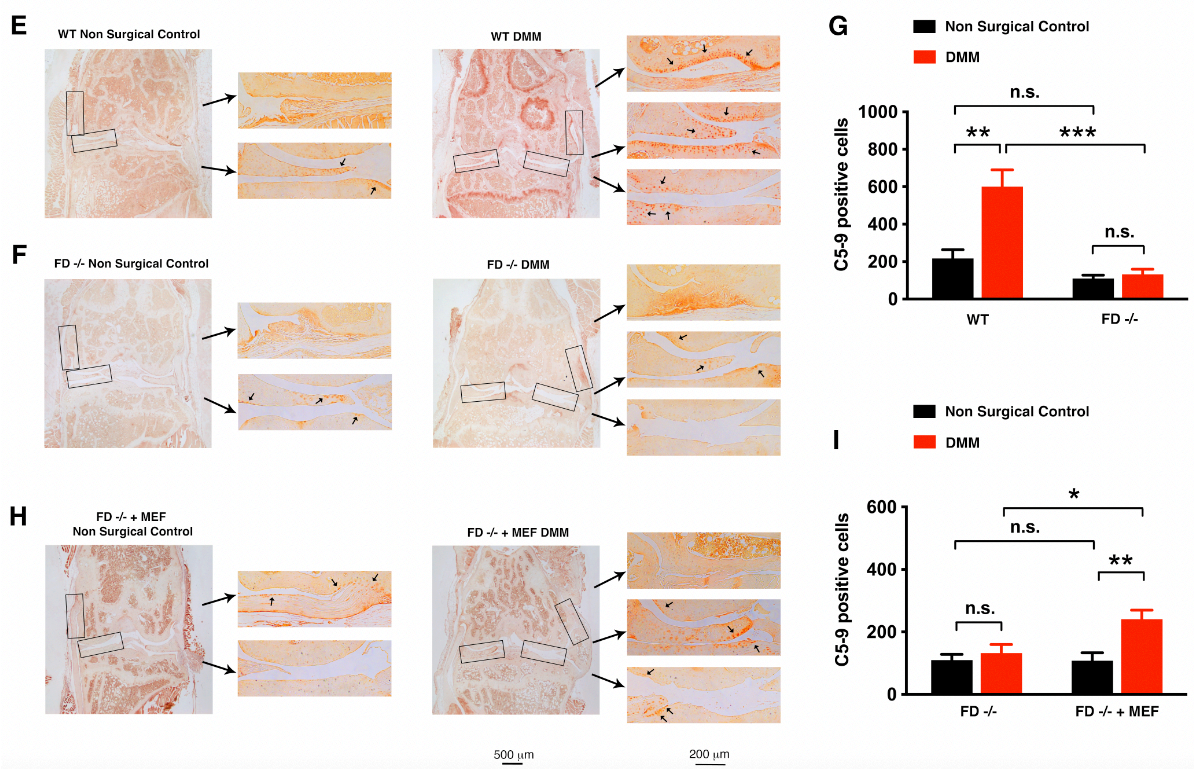

E
WT Non Surgical Control
WT DMM
F
FD -/- Non Surgical Control
FD -/- DMM
G
Non Surgical Control
DMM
C5-9 positive cells
1000
800
600
400
200
0
n.s.
**
***
n.s.
WT
FD -/-
H
FD -/- + MEF
Non Surgical Control
FD -/- + MEF DMM
I
Non Surgical Control
DMM
C5-9 positive cells
600
400
200
0
*
n.s.
**
n.s.
FD -/-
FD -/- + MEF
500 μm
200 μm

**Fig. S5. Assessment of complement activation in DMM joints using Immunohistochemistry (IHC).** (A) IHC staining for C3 in HFD WT HFD FD$^{-/-}$ and (B) quantification of C3 positive cells. (C) HFD FD$^{-/-}$ + MEF and (D) quantification of HFD FD$^{-/-}$ compared to HFD FD$^{-/-}$ + MEF. DMM limbs are demonstrated in red, non-surgical contralateral limbs are shown in black. C5-9 staining for (E) WT and (F) FD$^{-/-}$ and (G) quantification. (H) HFD FD$^{-/-}$ + MEF and (I) quantification of HFD FD$^{-/-}$ compared to HFD FD$^{-/-}$ + MEF. Each graph indicates n=5-8 samples/group. Comparisons were evaluated using 2-way ANOVA with x posthoc testing $p<0.05$). * Indicates $p<0.05$, ** indicates $p<0.01$. Black – contralateral limb, Red – DMM limb.

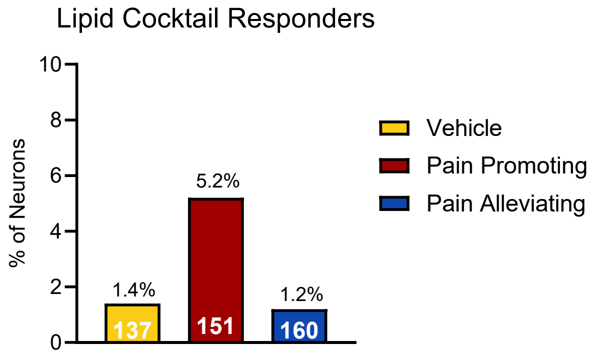


**Fig. S6. Cocktail application does not acutely activate DRG neurons.** Direct application of lipid cocktails during calcium imaging resulted in minimal acute calcium responses in DRG neurons. The percentage of responding neurons was low for both cocktails and was not significantly different from vehicle treatment.

**Table S1. Differentially regulated pathways across chow- and HFD-fed DMM and naïve mice identified by median metabolite intensity heatmap analysis.** Clusters defined in Fig. 1H. All pathways listed were false discover rate (FDR)-corrected.

| Cluster | Pathway | Pathway total | Hits.total | Hits.sig | FDR P-value |
|---|---|---|---|---|---|
| 1 | Ascorbate and aldarate metabolism | 10 | 6 | 4 | 0.008 |
| | Tryptophan metabolism | 41 | 23 | 8 | 0.009 |
| | Purine metabolism | 71 | 19 | 7 | 0.009 |
| | Citrate cycle (TCA cycle) | 20 | 11 | 5 | 0.009 |
| | Pentose and glucuronate interconversions | 19 | 7 | 3 | 0.019 |
| | Lysine degradation | 30 | 12 | 4 | 0.02 |
| | Drug metabolism - other enzymes | 38 | 8 | 3 | 0.024 |
| | Glyoxylate and dicarboxylate metabolism | 32 | 14 | 4 | 0.029 |
| | Alanine, aspartate and glutamate metabolism | 28 | 20 | 5 | 0.032 |
| | Amino sugar and nucleotide sugar metabolism | 42 | 20 | 5 | 0.032 |
| | Metabolism of xenobiotics by cytochrome P450 | 64 | 20 | 5 | 0.032 |
| | Butanoate metabolism | 15 | 4 | 2 | 0.034 |
| | Terpenoid backbone biosynthesis | 18 | 5 | 2 | 0.045 |
| | Drug metabolism - cytochrome P450 | 27 | 17 | 4 | 0.047 |
| 2 | Glutathione metabolism | 28 | 9 | 7 | 0.042 |
| | Valine, leucine and isoleucine biosynthesis | 8 | 4 | 4 | 0.045 |
| | Linoleic acid metabolism | 5 | 4 | 4 | 0.045 |
| | Lysine degradation | 30 | 12 | 7 | 0.047 |
| | Arginine and proline metabolism | 36 | 18 | 9 | 0.049 |
| 3 | No significant pathways | | | | |
| 4 | Biosynthesis of unsaturated fatty acids | 36 | 10 | 6 | 0.02 |
| | Lipoic acid metabolism | 28 | 5 | 4 | 0.021 |
| | alpha-Linolenic acid metabolism | 13 | 6 | 3 | 0.046 |
| 5 | Valine, leucine and isoleucine biosynthesis | 8 | 4 | 4 | 0.039 |
| | Purine metabolism | 71 | 19 | 8 | 0.039 |
| | Glycerophospholipid metabolism | 36 | 6 | 4 | 0.04 |
| | D-Amino acid metabolism | 15 | 11 | 5 | 0.041 |
| | Arginine and proline metabolism | 36 | 18 | 6 | 0.043 |
| | Glutathione metabolism | 28 | 9 | 4 | 0.044 |
| | Ubiquinone and other terpenoid-quinone biosynthesis | 18 | 6 | 3 | 0.048 |
| | Tryptophan metabolism | 41 | 23 | 6 | 0.049 |
| 6 | Biosynthesis of unsaturated fatty acids | 36 | 10 | 7 | 0.044 |
| | Retinol metabolism | 16 | 11 | 7 | 0.045 |
| 7 | No significant pathways | | | | |

**Table S2. Hallmark pathway enrichment analysis of differentially expressed serum proteins in HFD vs chow-fed DMM mice.** Differentially expressed proteins between DMM animals were identified from volcano plot analysis – as seen in Fig. 1N – then analyzed using iPathway to identify enriched hallmark pathways. Differentially expressed and all hits, p-values, and FDR-corrected p-values are reported for each hallmark signature.

| Hallmark Signature | Count Differentially Expressed | Count All | P-Value | FDR P-value |
|---|---|---|---|---|
| Coagulation | 26 | 132 | 2.18E-07 | 1.05E-05 |
| Glycolysis | 32 | 193 | 5.43E-07 | 1.30E-05 |
| Reactive Oxygen Species Pathway | 13 | 46 | 4.274E-06 | 6.84E-05 |
| Heme Metabolism | 29 | 185 | 6.195E-06 | 7.43E-05 |
| Xenobiotic Metabolism | 27 | 184 | 4.49E-05 | 4.31E-04 |
| MTORC1 Signaling | 27 | 192 | 9.58E-05 | 7.67E-04 |
| Adipogenesis | 26 | 198 | 3.99E-04 | 0.003 |
| Complement | 24 | 183 | 6.82E-04 | 0.004 |
| Fatty Acid Metabolism | 20 | 142 | 7.75E-04 | 0.004 |
| MYC Targets V1 | 24 | 192 | 0.001 | 0.007 |
| Epithelial Mesenchymal Transition | 24 | 194 | 0.002 | 0.007 |
| Hypoxia | 23 | 194 | 0.003 | 0.0132 |

**Table S3. Hallmark pathway enrichment analysis of differentially expressed serum proteins between HFD and chow-fed naïve mice.** Differentially expressed proteins between naïve animals were identified from volcano plot analysis – as seen in Fig. S1E **–** then analyzed using iPathway to identify enriched hallmark pathways. Differentially expressed and all hits, p-values, and FDR-corrected p-values are reported for each hallmark signature.

| Hallmark Signature | Count Differentially Expressed | Count All | P-Value | FDR P-value |
|---|---|---|---|---|
| MYC Targets V1 | 40 | 192 | 2.322E-9 | 1.12E-7 |
| MTORC1 Signaling | 37 | 192 | 8.74E-8 | 2.01E-06 |
| Reactive Oxygen Species Pathway | 16 | 46 | 1.25E-7 | 2.0E-06 |
| Xenobiotic Metabolism | 35 | 184 | 2.81E-7 | 3.37E-06 |
| Coagulation | 27 | 132 | 1.63E-06 | 1.56E-05 |
| Glycolysis | 32 | 193 | 2.00E-05 | 1.60E-04 |
| Heme Metabolism | 29 | 185 | 1.42E-04 | 9.71E-04 |
| Fatty Acid Metabolism | 22 | 142 | 0.001 | 0.006 |
| PI3K AKT MTOR Signaling | 17 | 104 | 0.002 | 0.012 |
| Myogenesis | 26 | 196 | 0.004 | 0.019 |
| Complement | 24 | 183 | 0.006 | 0.025 |
| Epithelial Mesenchymal Transition | 25 | 194 | 0.007 | 0.025 |
| Hypoxia | 25 | 194 | 0.007 | 0.025 |

**Table S4. Differentially regulated pathways across IDEA participants by change in WOMAC pain scores over 18 months of weight loss intervention identified by median metabolite intensity heatmap analysis.** Clusters defined in Fig. 4G. All pathways listed were FDR-corrected.

| Cluster | Pathway | Pathway total | Hits.total | Hits.sig | FDR p-value |
|---|---|---|---|---|---|
| 1 | Butanoate metabolism | 34 | 6 | 4 | 0.002 |
| | Propanoate metabolism | 31 | 4 | 3 | 0.003 |
| | Vitamin E metabolism | 54 | 5 | 3 | 0.004 |
| | Pyrimidine metabolism | 70 | 14 | 5 | 0.004 |
| | Tryptophan metabolism | 94 | 15 | 5 | 0.004 |
| | Purine metabolism | 80 | 15 | 5 | 0.004 |
| | Glycine, serine, alanine and threonine metabolism | 88 | 11 | 4 | 0.005 |
| | Glycosphingolipid biosynthesis - lactoseries | 14 | 2 | 2 | 0.005 |
| | Blood Group Biosynthesis | 44 | 2 | 2 | 0.005 |
| | Glycosphingolipid biosynthesis - neolactoseries | 16 | 2 | 2 | 0.005 |
| | Beta-Alanine metabolism | 20 | 2 | 2 | 0.005 |
| | Lysine metabolism | 52 | 13 | 4 | 0.008 |
| | Vitamin B3 (nicotinate and nicotinamide) metabolism | 28 | 8 | 3 | 0.008 |
| | Valine, leucine and isoleucine degradation | 65 | 3 | 2 | 0.008 |
| | Glycolysis and Gluconeogenesis | 49 | 9 | 3 | 0.011 |
| | N-Glycan biosynthesis | 48 | 4 | 2 | 0.013 |
| | Glycosphingolipid biosynthesis - globoseries | 16 | 4 | 2 | 0.013 |
| 2 | Vitamin D3 (cholecalciferol) metabolism | 16 | 5 | 2 | 0.01 |
| | Xenobiotics metabolism | 110 | 6 | 2 | 0.014 |
| | Urea cycle/amino group metabolism | 85 | 19 | 3 | 0.032 |
| | Fatty acid oxidation | 35 | 11 | 2 | 0.04 |
| | Fatty acid activation | 74 | 12 | 2 | 0.046 |
| 3 | Saturated fatty acids beta-oxidation | 36 | 3 | 2 | 0.013 |
| | Carnitine shuttle | 72 | 28 | 6 | 0.022 |
| | C21-steroid hormone biosynthesis and metabolism | 112 | 5 | 2 | 0.026 |
| | Phytanic acid peroxisomal oxidation | 34 | 5 | 2 | 0.026 |
| | Linoleic acid metabolism | 46 | 6 | 2 | 0.034 |
| | Xenobiotics metabolism | 110 | 6 | 2 | 0.034 |
| 4 | Glycerophospholipid metabolism | 156 | 17 | 8 | 0.004 |
| | Vitamin E metabolism | 54 | 5 | 3 | 0.009 |
| | Beta-Alanine metabolism | 20 | 2 | 2 | 0.012 |
| | Leukotriene metabolism | 92 | 9 | 4 | 0.012 |
| | Di-unsaturated fatty acid beta-oxidation | 26 | 6 | 3 | 0.014 |
| | Caffeine metabolism | 11 | 3 | 2 | 0.02 |
| | Pyruvate Metabolism | 20 | 4 | 2 | 0.032 |
| | Glycolysis and Gluconeogenesis | 49 | 9 | 3 | 0.041 |
| | Fructose and mannose metabolism | 33 | 9 | 3 | 0.041 |
| | Mono-unsaturated fatty acid beta-oxidation | 19 | 5 | 2 | 0.048 |
| | Ascorbate (Vitamin C) and Aldarate Metabolism | 29 | 5 | 2 | 0.048 |
| 5 | Biopterin metabolism | 22 | 3 | 2 | 0.007 |
| | Pyrimidine metabolism | 70 | 14 | 3 | 0.024 |
| | Tryptophan metabolism | 94 | 15 | 3 | 0.029 |
| 6 | Drug metabolism - cytochrome P450 | 53 | 11 | 6 | 0.002 |
| | Vitamin B6 (pyridoxine) metabolism | 11 | 3 | 3 | 0.003 |
| | Putative anti-Inflammatory metabolites formation from EPA | 27 | 2 | 2 | 0.007 |
| | Phosphatidylinositol phosphate metabolism | 59 | 6 | 3 | 0.007 |
| | Valine, leucine and isoleucine degradation | 65 | 3 | 2 | 0.012 |
| | Dynorphin metabolism | 8 | 3 | 2 | 0.012 |
| | Drug metabolism - other enzymes | 31 | 3 | 2 | 0.012 |
| | Arachidonic acid metabolism | 95 | 4 | 2 | 0.019 |
| | Glycosphingolipid metabolism | 67 | 14 | 4 | 0.019 |
| | Tryptophan metabolism | 94 | 15 | 4 | 0.025 |
| | Alkaloid biosynthesis II | 10 | 5 | 2 | 0.028 |
| | Glycine, serine, alanine and threonine metabolism | 88 | 11 | 3 | 0.032 |
| | Butanoate metabolism | 34 | 6 | 2 | 0.038 |
| 7 | Porphyrin metabolism | 43 | 4 | 3 | 0.007 |
| | N-Glycan Degradation | 16 | 4 | 3 | 0.007 |
| | Histidine metabolism | 33 | 7 | 4 | 0.007 |
| | Pentose phosphate pathway | 37 | 7 | 4 | 0.007 |
| | Urea cycle/amino group metabolism | 85 | 19 | 7 | 0.013 |
| | Aminosugars metabolism | 69 | 10 | 4 | 0.02 |
| | Biopterin metabolism | 22 | 3 | 2 | 0.024 |
| | Glutamate metabolism | 15 | 3 | 2 | 0.024 |
| | Prostaglandin formation from arachidonate | 78 | 3 | 2 | 0.024 |
| | Glycine, serine, alanine and threonine metabolism | 88 | 11 | 4 | 0.027 |
| | Tryptophan metabolism | 94 | 15 | 5 | 0.028 |
| | Vitamin B3 (nicotinate and nicotinamide) metabolism | 28 | 8 | 3 | 0.035 |
| | N-Glycan biosynthesis | 48 | 4 | 2 | 0.038 |
| | Propanoate metabolism | 31 | 4 | 2 | 0.038 |
| | Glutathione Metabolism | 19 | 4 | 2 | 0.038 |
| | Starch and Sucrose Metabolism | 33 | 4 | 2 | 0.038 |

**Table S5. Pain-promoting and pain-alleviating cocktails generated from HFD FD$^{-/-}$ and MEF-treated HFD FD$^{-/-}$ serum profiles and pressure-pain hyperalgesia outcomes.** For each individual lipid within the pain-promoting and pain-alleviating cocktail its regulation pattern, the Cayman product number, and working concentration are reported.

| **Pain Promoting Cocktail** | | | |
|---|---|---|---|
| **Lipid** | **Regulation** | **Cayman Product Number** | **Working Concentration** |
| 5,6 DiHETE | FD-/- | 10467 | 10nM |
| U-46619 (Thromboxane B2 sub) | FD-/- | U-46619 | 10nM |
| 13,14-dihydro-15-keto PGF2a | FD-/- | 10007793 | 1nM |
| **Pain Alleviation Cocktail** | | | |
| **Lipid** | **Regulation** | **Cayman Product Number** | **Working Concentration** |
| 12,13 EpOME | FD-/- + MEF | 52450 | 1nM |
| 4-HDHA | FD-/- + MEF | 33200 | 1nM |
| DHA (docosahexaenoic acid) | FD-/- + MEF | 90310 | 1uM |
| 13-OxoODE | FD-/- + MEF | 38620 | 1nM |
| Linoleic Acid | FD-/- + MEF | 90150 | 1uM |
| EPA (eicosapentaenoic acid) | FD-/- + MEF | 90110 | 1uM |
| 10-Nitrolinoleate | FD-/- + MEF | 10037 | 10nM |
| 12,13 DiHOME | FD-/- + MEF | 10009832 | 1nM |
| DPA (docosapentaenoic acid) | FD-/- + MEF | 90165 | 1uM |
| 9,10 EpOME | FD-/- + MEF | 52400 | 1nM |

**Table S6. Statistical results for acute, short-, and long-term treatment of DRGs with pain-promoting and pain-alleviating cocktails.** Summary of statistical analyses for calcium imaging experiments assessing capsaicin and KCl-evoked responses in human DRGs following cocktail treatment. Comparisons include proportions of responsive neurons, response magnitude, latency, and area under the curve (AUC) across acute, short-term, and long-term treatment exposures.

| | | | | | |
|---|---|---|---|---|---|
| **24-hour Treatment Data** | | | | | |
| **Proportion of Capsaicin Responders** | | | | | |
| **Group** | **Responders** | **Non-responders** | **% Responders** | **Fishers Exact Test Compared with Vehicle** | **Fishers Exact Test Compared with Pain Promoting** |
| Vehicle | 100 | 61 | 62% | X | p<0.01 |
| Pain Promoting | 78 | 116 | 40% | p<0.01 | X |
| Pain Alleviating | 89 | 109 | 45% | p<0.01 | p=1.0 |
| **Magniude of Capsaicin Responders** | | | | | |
| **Group** | **Mean** | **SD** | **# of Neurons** | **One-way Anova** | **Tukey Post Hoc Test** |
| Vehicle | 93.01 | 58.95 | 100 | | |
| Pain Promoting | 92.33 | 56.25 | 78 | $F_{2,264}$ = 1.01, p=0.37 | X |
| Pain Alleviating | 82.75 | 45.89 | 89 | | |
| **Latency to Capsaicin Response** | | | | | |
| **Group** | **Mean** | **SD** | **# of Neurons** | **One-way Anova** | **Tukey Post Hoc Test** |
| Vehicle | 51.56 | 25.74 | 100 | | |
| Pain Promoting | 56.63 | 25.30 | 78 | $F_{2,264}$ = 1.10, p=0.33 | X |
| Pain Alleviating | 55.92 | 24.35 | 89 | | |
| **AUC of Capsaicin Response** | | | | | |
| **Group** | **Mean** | **SD** | **# of Neurons** | **One-way Anova** | **Tukey Post Hoc Test** |
| Vehicle | 21.63 | 14.95 | 100 | | |
| Pain Promoting | 22.76 | 16.91 | 78 | $F_{2,264}$ = 0.73, p=0.48 | X |
| Pain Alleviating | 19.92 | 14.54 | 89 | | |
| **Magnitude of KCl Response** | | | | | |
| **Group** | **Mean** | **SD** | **# of Neurons** | **One-way Anova** | **Tukey Post Hoc Test** |
| Vehicle | 65.94 | 37.66 | 61 | | |
| Pain Promoting | 74.00 | 34.15 | 116 | $F_{2,283}$ = 1.05, p=0.35 | X |
| Pain Alleviating | 69.71 | 37.60 | 109 | | |
| **Latency to KCl Response** | | | | | |
| **Group** | **Mean** | **SD** | **# of Neurons** | **One-way Anova** | **Tukey Post Hoc Test** |
| Vehicle | 19.62 | 7.70 | 61 | | Vehicle vs Promoting: p<0.01 |
| Pain Promoting | 16.07 | 5.30 | 116 | $F_{2,283}$ = 12.19, p<0.01 | Promoting vs Alleviating: p<0.01 |
| Pain Alleviating | 19.80 | 6.06 | 109 | | Vehicle vs Alleviating: p=0.98 |
| **30 Minute Treatment Data** | | | | | |
| **Proportion of Capsaicin Responders** | | | | | |
| **Group** | **Responders** | **Non-responders** | **% Responders** | **Fishers Exact Test Compared with Vehicle** | **Fishers Exact Test Compared with Pain Promoting** |
| Vehicle | 56 | 75 | 42% | X | p=0.04 |
| Pain Promoting | 91 | 67 | 57% | p=0.04 | X |
| Pain Alleviating | 58 | 46 | 55% | p=0.15 | p=1.0 |
| **Magnitude of Capsaicin Responders** | | | | | |
| **Group** | **Mean** | **SD** | **# of Neurons** | **One-way Anova** | **Tukey Post Hoc Test** |
| Vehicle | 82.21 | 48.77 | 56 | | |
| Pain Promoting | 100.10 | 52.91 | 91 | $F_{2,202}$ = 2.48, p=0.09 | X |
| Pain Alleviating | 86.42 | 51.44 | 58 | | |
| **Latency to Capsaicin Response** | | | | | |
| **Group** | **Mean** | **SD** | **# of Neurons** | **One-way Anova** | **Tukey Post Hoc Test** |
| Vehicle | 67.67 | 25.23 | 56 | | Vehicle vs Promoting: p<0.01 |
| Pain Promoting | 53.7 | 25.50 | 91 | $F_{2,202}$ = 5.64, p<0.01 | Promoting vs Alleviating: p=0.14 |
| Pain Alleviating | 61.59 | 23.96 | 58 | | Vehicle vs Alleviating: p=0.40 |
| **AUC of Capsaicin Response** | | | | | |
| **Group** | **Mean** | **SD** | **# of Neurons** | **One-way Anova** | **Tukey Post Hoc Test** |
| Vehicle | 17.58 | 15.00 | 56 | | Vehicle vs Promoting: p=0.01 |
| Pain Promoting | 25.44 | 17.64 | 91 | $F_{2,202}$ = 4.23, p=0.02 | Promoting vs Alleviating: p=0.18 |
| Pain Alleviating | 20.54 | 15.87 | 58 | | Vehicle vs Alleviating: p=0.60 |
| **Magnitude of KCl Response** | | | | | |
| **Group** | **Mean** | **SD** | **# of Neurons** | **One-way Anova** | **Tukey Post Hoc Test** |
| Vehicle | 64.41 | 28.90 | 75 | | |
| Pain Promoting | 64.99 | 31.97 | 67 | $F_{2,184}$ = 0.18, p=0.83 | X |
| Pain Alleviating | 67.79 | 31.13 | 46 | | |
| **Latency to KCl Response** | | | | | |
| **Group** | **Mean** | **SD** | **# of Neurons** | **One-way Anova** | **Tukey Post Hoc Test** |
| Vehicle | 18.28 | 3.94 | 75 | | |
| Pain Promoting | 18.19 | 5.71 | 67 | $F_{2,184}$ = 0.24, p=0.83 | X |
| Pain Alleviating | 17.70 | 3.92 | 46 | | |
| **Acute Lipid Treatment** | | | | | |
| **Group** | **Responders** | **Non-responders** | **% Responders** | **Fishers Exact Test Compared with Vehicle** | **Fishers Exact Test Compared with Pain Promoting** |
| Vehicle | 2 | 135 | 1.4% | X | p=0.32 |
| Pain Promoting | 8 | 143 | 5.2% | p=0.32 | X |
| Pain Alleviating | 2 | 158 | 1.2% | p=1.0 | p=0.17 |

**Table S7. Organ donor demographic data.** Human DRGs were recovered from 8 organ donors, including both males and females, where the mean age was 36.4 years.

| **Sex** | **N** | **Mean Age (years)** |
|---|---|---|
| Male | 5 | 34.2 |
| Female | 3 | 40 |
| **Total** | 8 | 36.4 |
